# Closed-loop optogenetic perturbation of macaque oculomotor cerebellum: evidence for an internal saccade model

**DOI:** 10.1101/2023.06.22.546199

**Authors:** Robijanto Soetedjo, Gregory D. Horwitz

**Affiliations:** Department of Physiology and Biophysics, University of Washington, Seattle, WA 98195, USA; Washington National Primate Research Center, University of Washington, Seattle, WA 98195, USA

## Abstract

Internal models are essential for the production of accurate movements. The accuracy of saccadic eye movements is thought to be mediated by an internal model of oculomotor mechanics encoded in the cerebellum. The cerebellum may also be part of a feedback loop that predicts the displacement of the eyes and compares it to the desired displacement in real time to ensure that saccades land on target. To investigate the role of the cerebellum in these two aspects of saccade production, we delivered saccade-triggered light pulses to channelrhodopsin-2-expressing Purkinje cells in the oculomotor vermis (OMV) of two macaque monkeys. Light pulses delivered during the acceleration phase of ipsiversive saccades slowed the deceleration phase. The long latency of these effects and their scaling with light pulse duration are consistent with an integration of neural signals at or downstream of the stimulation site. In contrast, light pulses delivered during contraversive saccades reduced saccade velocity at short latency and were followed by a compensatory reacceleration which caused gaze to land near or on the target. We conclude that the contribution of the OMV to saccade production depends on saccade direction; the ipsilateral OMV is part of a forward model that predicts eye displacement, whereas the contralateral OMV is part of an inverse model that creates the force required to move the eyes with optimal peak velocity for the intended displacement.

**Significance Statement:** Theory and experiment suggest that saccade production involves an internal model of oculomotor mechanics that resides inside the cerebellum. How cerebellar neurons implement this model is poorly understood. To illuminate this issue, we stimulated Purkinje cells in the oculomotor vermis (OMV) optogenetically during saccades and examined the resultant movement deviations. Stimulation of the contralateral OMV affected saccade dynamics at short latency, suggesting that the contralateral OMV is part of the feedforward pathway that produces the saccade motor command. In contrast, perturbation of the ipsilateral OMV affected saccade dynamics at longer latency, prolonging the saccade deceleration phase and leading to hypermetria. These effects are consistent with perturbation of the eye displacement integrator in the feedback loop of the saccade generator.

## Introduction

The generation of accurate saccades requires an intact oculomotor vermis (OMV, lobules VIc and VII) (Takagi et al., 1998; Barash et al., 1999). The OMV receives mossy fiber inputs from pontine areas critical for producing saccades (Yamada and Noda, 1987, Fig. 1A): the nucleus reticularis tegmenti pontis (NRTP), which is thought to relay the saccade command from the superior colliculus (SC) (Crandall and Keller, 1985), and saccade premotor burst neurons in the saccade burst generator (SBG). Purkinje (P-) cells in the OMV project to the ipsilateral caudal fastigial nucleus (cFN) which, in turn, projects to the premotor burst neurons (Noda et al., 1990). This close connectivity predicts that P-cell activity is closely related to saccadic eye movements (Herzfeld et al., 2015), and consistent with this role, electrical stimulation of the OMV drives the eyes ipsilaterally at short latency (Fujikado and Noda, 1987; Krauzlis and Miles, 1998). These observations narrow the possible roles of the OMV in eye movement control but do not identify how OMV P-cells map onto components of quantitative models of saccade control (Fuchs et al., 1985).

**Figure 1.**
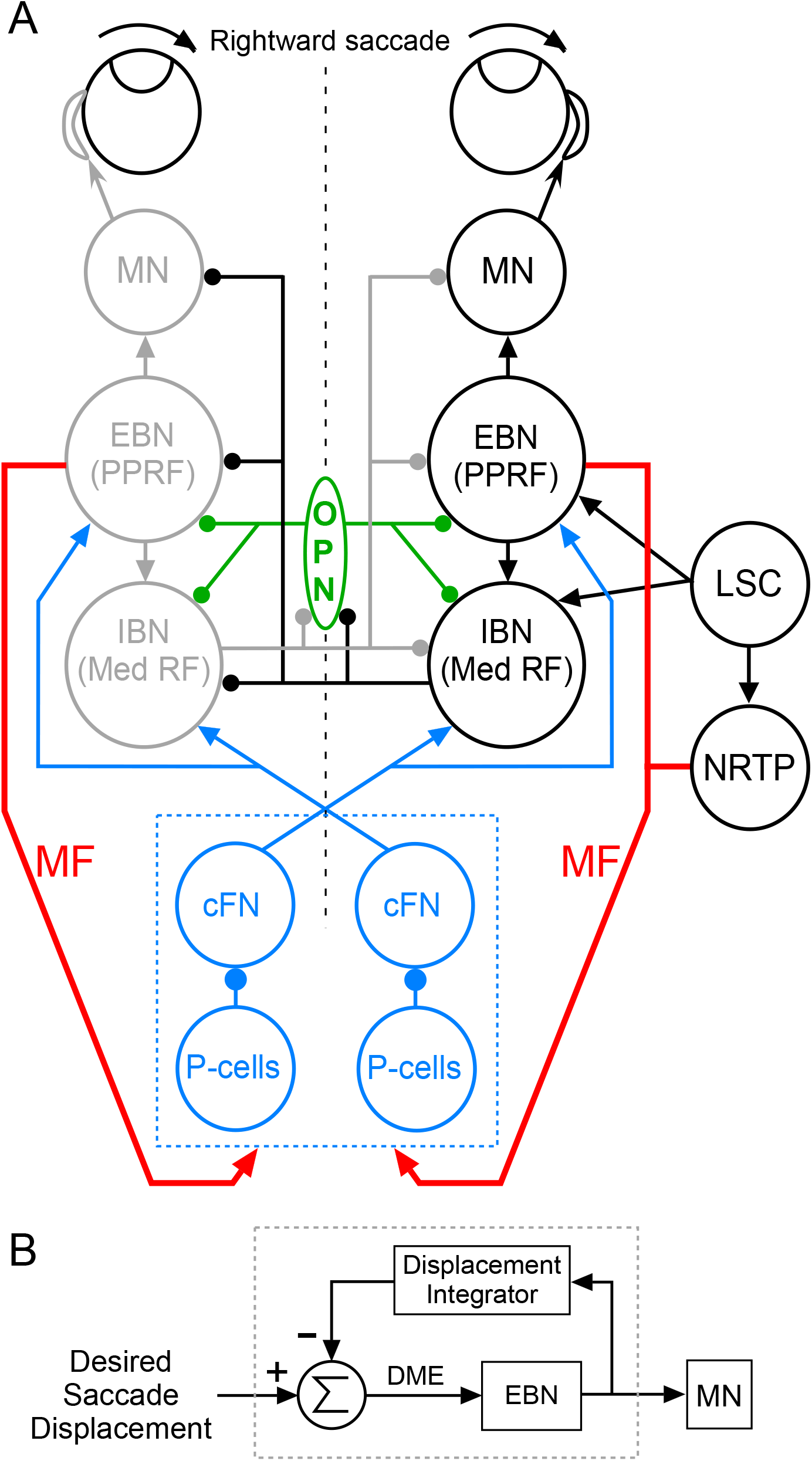
Simplified block diagrams of the horizontal saccade generator (A) and the local negative feedback model of the saccade burst generator (B, gray dashed box). During fixation, omnipause neurons (OPN) inhibit both excitatory burst neurons (EBN) in the paramedian pontine reticular formation (PPRF) and inhibitory burst neurons (IBN) in the medullary reticular formation (Med RF). To produce a rightward saccade, the left superior colliculus (LSC) sends a desired saccade displacement command to the right EBNs and IBNs. The right IBNs inhibit the OPNs, thereby allowing both right EBNs and IBNs to fire a burst. This burst drives the abducens motoneurons (MN) to rotate the eyes rightward. The burst of right IBNs inhibit left MNs, EBNs and IBNs. The nucleus reticularis tegmenti pontis (NRTP), which receives input from the SC, projects as mossy fibers (MF, red arrows) to the oculomotor vermis and caudal fastigial nucleus (cFN). The cFN projects to contralateral EBNs and IBNs. Purkinje cells (P-cells) inhibit neurons in the cFN. Blue dashed box, cerebellum. DME, dynamic motor error.

According to an established local feedback model (Fig. 1B), a desired saccade displacement command drives excitatory burst neurons (EBN), which, in turn, activate motoneurons (MN) to move the eyes quickly to the intended target (Robinson, 1975; Jürgens et al., 1981). This feedforward pathway produces the force needed to achieve a velocity profile optimal for the intended displacement (Robinson, 1964; Harris and Wolpert, 2006).

A second component of the model is a feedback pathway that integrates an efference copy of the EBN command to predict the displacement of the eyes. When the difference between the predicted and desired displacement (the dynamic motor error [DME], Fig. 1B) reaches zero, the saccade stops. This model explains how saccade accuracy is maintained despite midflight perturbations (Keller et al., 1996; Xu-Wilson et al., 2011) or pharmacological slowing (Soetedjo et al., 2002; Barton et al., 2003).

Whether the OMV/cFN complex plays a role in the closed-loop control of saccades is unclear. The fact that the OMV/cFN complex receives a copy of the saccade command from the NRTP and a feedback signal from premotor neurons suggests that it could be part of the local feedback circuit. However, electrical stimulation experiments suggest that the cFN plays a role in modulating the saccade command in an open-loop fashion (Noda et al., 1991).

Resolving this uncertainty is difficult with electrical microstimulation, which excites multiple classes of neurons and passing fibers synchronously (Noda et al., 1988). Inactivation methods may keep neural circuits closer to their normal state but are difficult to implement with the speed and reversibility necessary to distinguish open-loop from closed-loop controllers.

To achieve fast, reversible inactivation of the cFN, we used optogenetics to activate OMV P-cells, which inhibit cFN neurons. Specifically, we drove channelrhodopsin-2 expression in OMV P-cells using the P-cell-specific L7 promoter and activated them with brief light pulses during visually guided saccades.

Effects of these light pulses depended on saccade direction. Contraversive saccades were slowed in midflight after a short latency, and this deceleration was followed by a compensatory reacceleration that brought the eyes to the target. Both effects are consistent with reduced drive in the feedforward pathway. In contrast, stimulation during ipsiversive saccades prolonged their deceleration, largely independently of when the stimulation was applied. These saccades were systematically hypermetric, and the amount of hypermetria scaled with light pulse duration. Both effects are consistent with a reduced drive at the input of the feedback integrator. We conclude that the role of the OMV in saccade control depends on the direction of the saccade: the contralateral OMV is part of the feedforward pathway whereas the ipsilateral OMV is part of the feedback integrator. This conclusion is supported by a detailed version of the saccade generator model (Fig. 13A) under which stimulation of the feedback and feedforward pathways produces perturbed eye movement trajectories that closely match those produced by actual brief activation of the ipsi-and contralateral OMV, respectively.

## Materials and Methods

All experiments were performed in accordance with the Guide for the Care and Use of Laboratory Animals and exceeded the minimum requirements of the Institute of Laboratory Animal Resources and the Association for Assessment and Accreditation of Laboratory Animal Care International. All the procedures were evaluated and approved by the Animal Care and Use Committee of the University of Washington.

### Animal subjects

Two juvenile male rhesus monkeys (*Macaca mulatta*), Sq and Tm, participated in this study. Sq, but not Tm, had participated in previous studies (El-Shamayleh et al., 2017; Soetedjo et al., 2019). Both animals underwent two surgeries; the first was to implant a scleral search eye coil (Judge et al., 1980) and head stabilization lugs. The second was to implant stainless steel recording chambers, one aimed at the superior colliculus (SC) and the other at the OMV. The OMV chamber was aimed straight down, 3 mm left of the midline and 14 mm posterior in Sq and 1.5 mm left of the midline and 15 mm posterior in Tm, relative to the stereotaxic ear-bar zero.

After the animals had recovered from the first surgery, they were trained to track a red laser dot (∼0.25° diameter) on a tangent screen 62 cm from their eyes. To obtain a dollop of fortified applesauce reward, the animals were required to fixate the dot for 1.2 – 2 sec, and after the target spot jumped, to shift their gaze to the target in <450 ms and remain within (± 2°) of the target for 1.5 sec. Once they performed this task reliably for ≥2 hours, the recording chambers were implanted.

### AAV vector Injection

An adeno-associated viral vector (AAV) carrying the channelrhodopsin-2 (ChR2) gene under the control of the P-cell-specific L7 promoter has been described previously (El-Shamayleh et al., 2017). This vector was injected into the OMV, which was identified on the basis of saccade-related activity (Fig. 2B), through an injectrode made of 30 G stainless steel hypodermic tubing insulated with polyimide and epoxylite (Kojima et al., 2021; Soetedjo and Kojima, 2022). Injection tracks were based on the presence of strong eye movement-related multiunit activity. (Soetedjo and Fuchs, 2006; Soetedjo et al., 2008; Kojima et al., 2010). Single units were isolated only rarely, and injections were made on the basis of saccade-related activity without regard to position of the functional midline. Sq received a total of 24 µL of AAV9-L7-ChR2-mCherry (1.41×10^13^ genomes/mL) across 4 tracks (∼0.75 mm apart) at up to 4 depths per track (1.5–2 µL at each depth over the course of 1 min with ≥ 2 min between injections) (El-Shamayleh et al., 2017). Tm received a total of 27 µL of AAV1-L7-ChR2- mCherry (1.53×10^13^ genomes/mL) along a single track at 7 depths (3–5 µL at each depth over the course of 2 min with ≥ 3 min between injections). Experiments began ≥ 6 weeks after injections.

**Figure 2.**
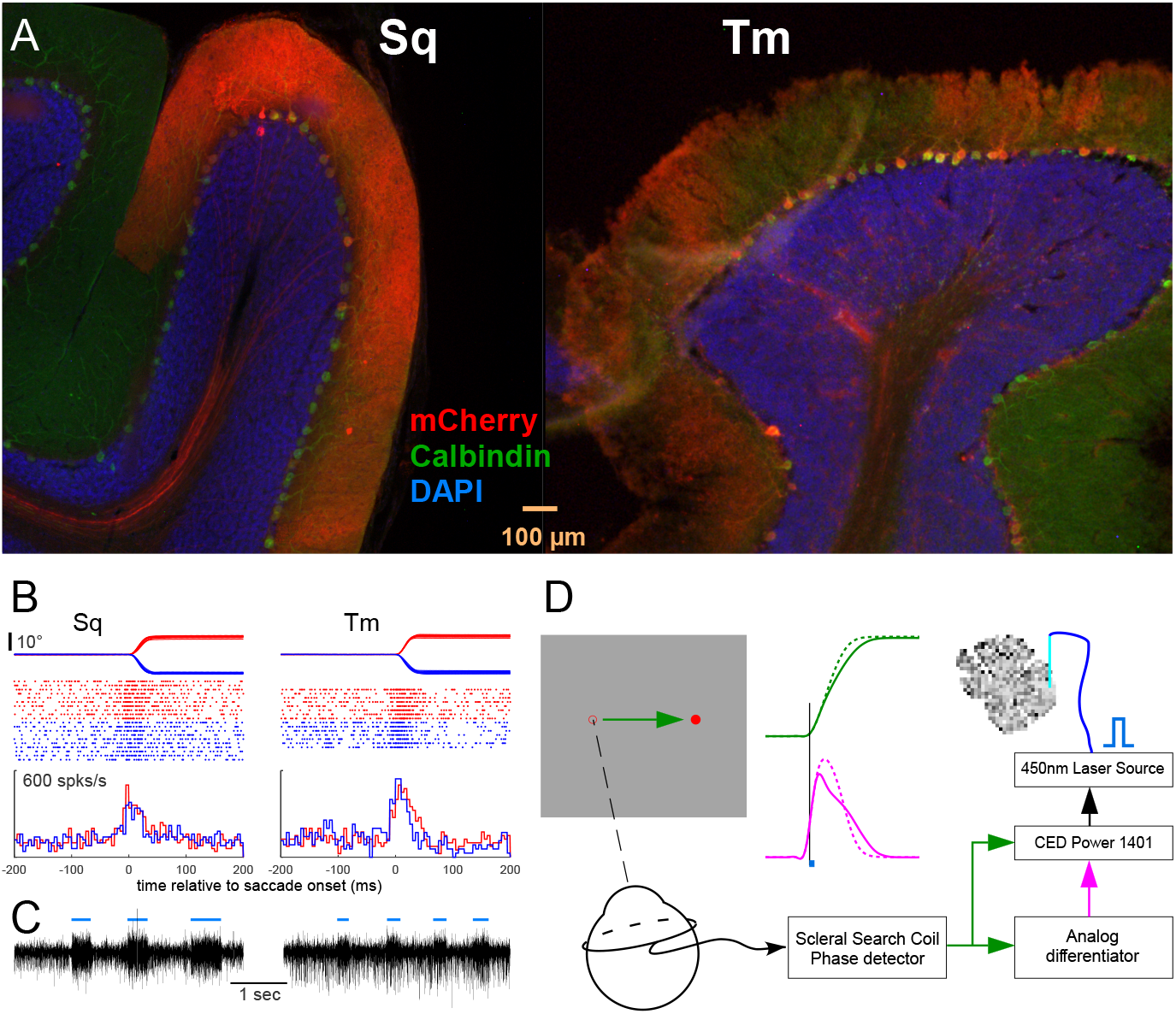
*A*, Transduction Pattern of AAV9–L7–ChR2–mCherry (Sq) and AAV1–L7–ChR2– mCherry (Tm) in cerebellar vermis lobule VIc. *B*, Multiunit activity during rightward (red) and leftward (blue) saccades. From top to bottom: eye position, spike rasters, and histograms. *C*, Analog multiunit trace during stimulation with blue laser light (blue bars, 20 mW). *D*, Systems for eye movement recording and light pulse delivery. Left panel shows the stepping target red dot on the screen and the scleral search coil. Middle panel shows normal eye position (solid green) and velocity (solid magenta) traces, and the effects of a light pulse (small blue bar, vertical line is light onset) on eye position and velocity (dashed curves).

### Histology

Histological methods have been described previously (El-Shamayleh et al., 2017). After the brain was fixed, sagittal sections (50 µm) of the cerebellum were cut and incubated with antibodies against the reporter protein mCherry (Clontech 632543, RRID: AB_2307319, 1:500) and against the P-cell marker calbindin (Swant CB38, RRID: AB_10000340). Sections were then stained with fluorescently tagged secondary antibodies (Thermo Fisher Scientific Cat# A10037, RRID: AB_2534013 and A32790, RRID:AB_2762833) and the nuclear stain, DAPI (Invitrogen Molecular Probes D-21490, 1:5000). They were then mounted onto glass slides and coverslipped. The expression of ChR2 was verified in the OMV of both animals (Fig. 2A).

### Saccade task

Sq and Tm were trained to make horizontal 24° saccades that started from a 12° eccentricity and crossed straight-ahead gaze. To reduce target predictability, approximately 30% of the steps were 6 or 18°. These catch trials were not accompanied by light pulses and were not analyzed. When a saccade was detected, the target spot was extinguished for 200 ms to reduce the possibility of any visually guided corrective movement following primary saccades. We did not find any signs of consistent corrective saccades up to ∼200 ms after saccade termination.

### Light activation

We stimulated ChR2-expressing P-cells with a custom built 450 nm light source based on an Osram PLTB450B laser diode (El-Shamayleh et al., 2017; Soetedjo and Kojima, 2022) coupled to a 200 µm, 0.39 NA (FT200EMT, Thorlabs, Newton, NJ, USA) optical fiber patch cable. To minimize tissue damage and increase light spread, the optical fiber was sharpened to an apical angle of 20°. A double-barrel cannula housed an Alpha-Omega glass-tungsten microelectrode (impedance: ∼100 kΩ at 1 kHz) and the optical fiber. The microelectrode was used to identify depths with strong saccade-related activity. The electrode and optical fiber were driven independently by an EPS micro-positioner (Alpha-Omega, Nazareth, Israel). During experiments, the tip of the fiber was 0.5–1 mm above the tip of the microelectrode. Experiments were conducted at sites at which multiunit activity was modulated by both saccades and optical stimulation (Figs. 2B and 2C).

We performed two types of experiment which differed in light pulse duration. In the first type, which was performed on both animals, the light pulse was triggered at one of 4 times relative to saccade initiation. The T0 light pulse was as early as possible (0 ms delay). The second, third and fourth trigger times occurred 10, 20, and 30 ms after T0 (T10, T20 and T30, respectively). T0 and T10 occurred during the saccade acceleration phase, T20 occurred near the time of peak velocity, and T30 occurred during the deceleration phase. In these experiments, light pulse duration was fixed at 30 ms for Sq and at 10 ms for Tm. The briefer pulses in Tm were motivated by the need for more precise estimates of effect latency, which differed for ipsilateral and contralateral saccades. To provide approximately equal light flux to each animal, the optical power of the laser was set to 60 mW for Sq and to 130 mW for Tm (Thorlabs PM100D power meter with S121C sensor). In the second experiment type, performed on Tm only, light pulses always began at T0 and had a duration drawn from four equally likely values: 1.25, 2.5, 5 and 10 ms.

The light pulse was triggered by saccade initiation (Fig. 2D). Saccade initiation was detected by a Power 1401 data acquisition and control system (Cambridge Electronic Design, UK) that monitored horizontal eye velocity continuously. When the eye velocity after a target step exceeded a 15°/s threshold, the Power 1401 randomly picked a trigger time and produced a TTL pulse. In the first experiment type, 4 trigger times were used (T0–T30), and the TTL pulse drove the laser directly. In the second type, the Power 1401 generated a TTL pulse at time T0 that triggered an Arduino microcontroller which randomly produced a 1.25, 2.5, 5 or 10 ms pulse that drove the laser. The delay introduced by the Arduino was ∼100 µs.

### General data analysis

All data were saved in Spike2 (Cambridge Electronic Design, UK) format, and analyses were performed in Spike2 and Matlab (Mathworks, USA). Eye and target position data were digitized at 1 kHz. Extracellular voltages and signals from a laser-detecting photodiode were digitized at 50 kHz. The on and off times of each laser pulse were extracted from the sampled photodiode output. Eye position traces were filtered forward and backward (Matlab “filtfilt” function, non-causal) with a 70 Hz, low pass, 61-coefficient Hamming window finite impulse response filter to reduce recording noise. A central derivative (Matlab “gradient” function) was applied successively to the filtered eye position signal to obtain eye velocity and then acceleration. After each target step, the program searched the subsequent 90–500 ms for an eye velocity >50 °/s and identified the first such movement, if it existed, as a saccade. Horizontal eye velocities that rose above, or fell below, a threshold of 15 °/s were considered the onset and termination of the saccade, respectively. The program identified the amplitude of the target step and measured the metrics of the saccades, as well as the occurrence times of light pulses. We discarded trials that were contaminated by blinks and those in which the horizontal or vertical eye position was ≥1.5° from the target when it stepped. These two criteria eliminated <10% of the trials, leaving a total of 1658 and 2856 for Sq and Tm, respectively (see Tables 1 and 2 as well as Fig. 11 legend for the number of trials in each stimulation condition).

**Table 1.**
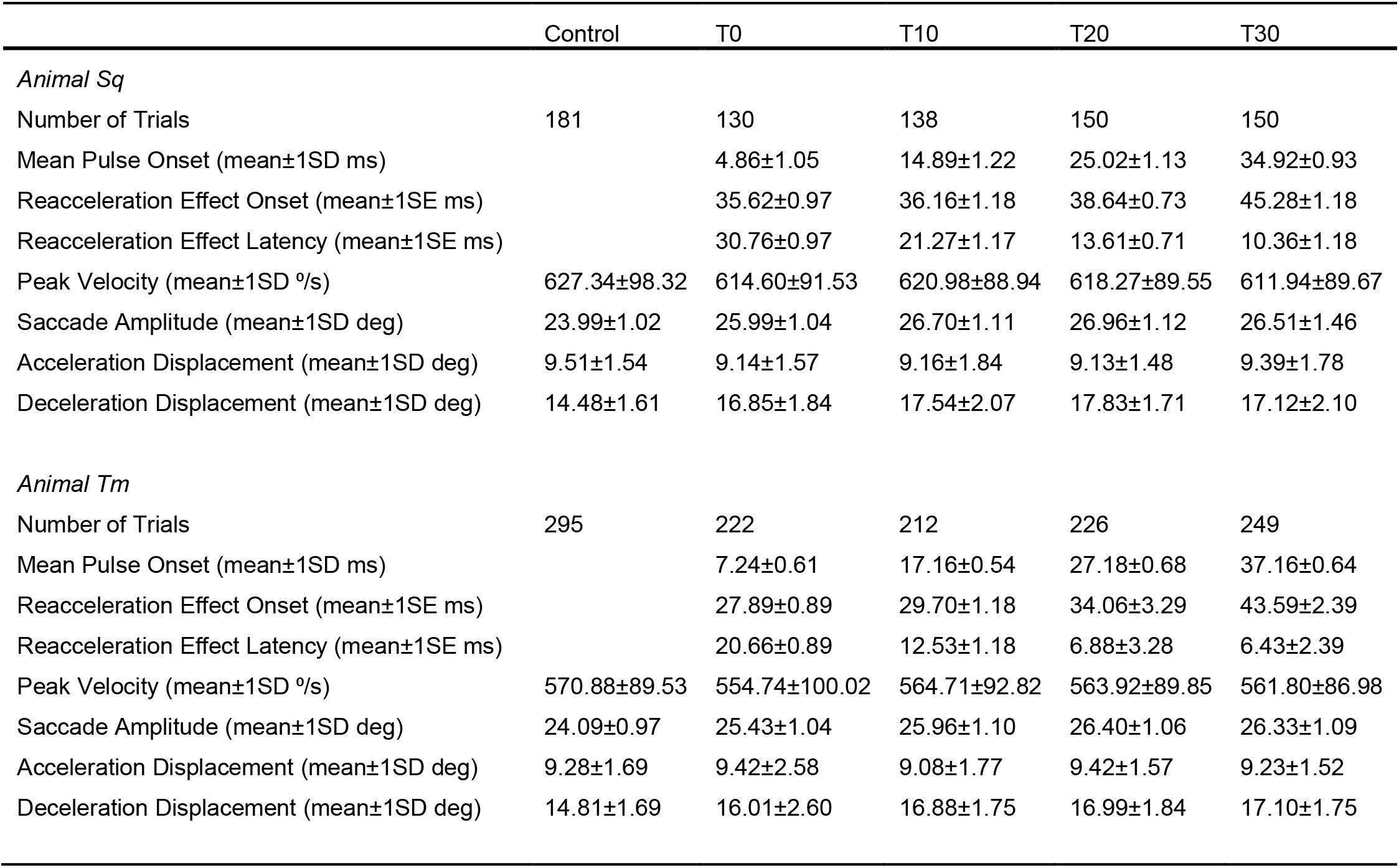
Summary of ipsiversive saccade metrics and optical activation timing.

**Table 2.**
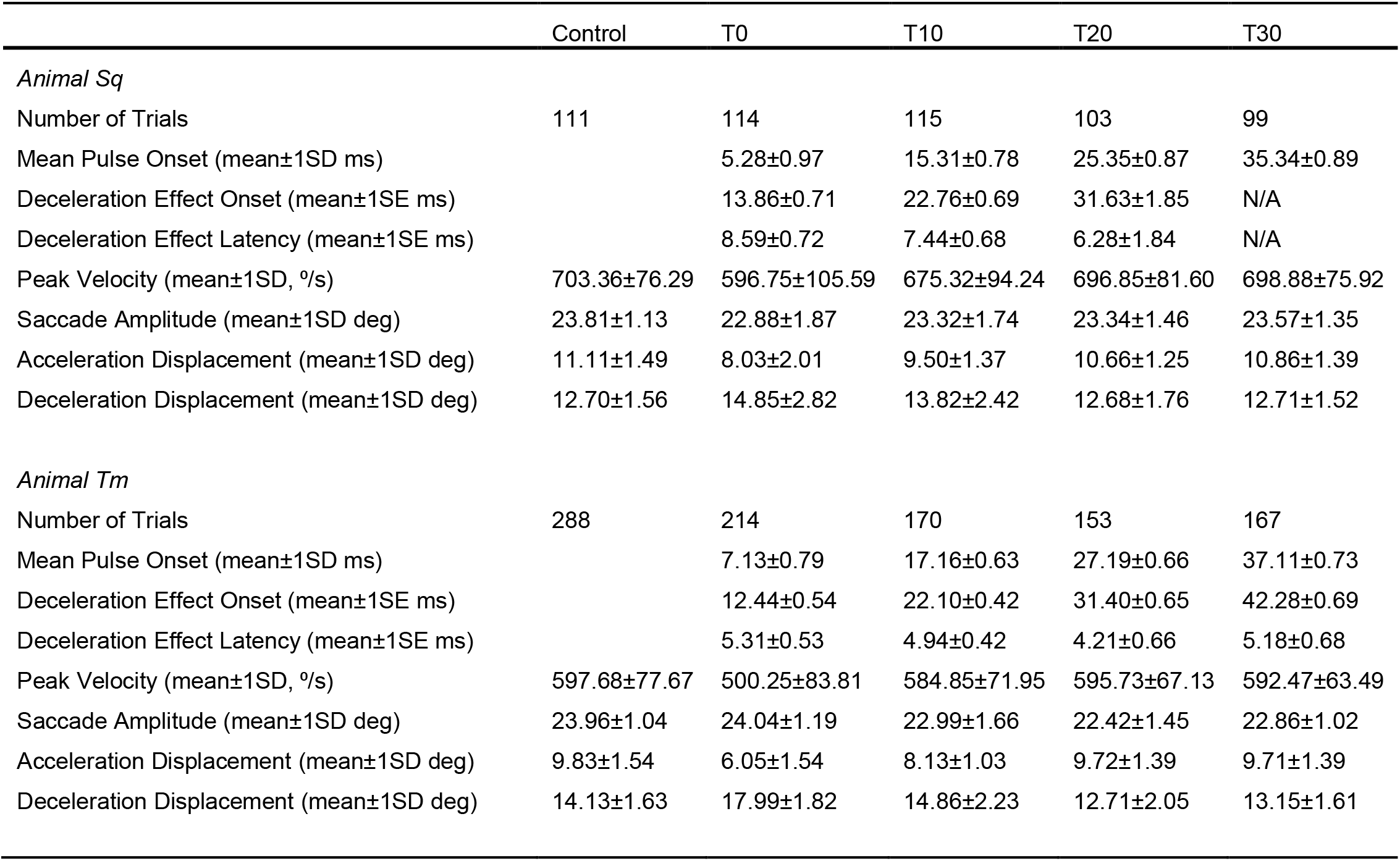
Summary of contraversive saccade metrics and optical activation timing.

### Determining optogenetic effect latency

To measure the latency of optogenetic perturbations of saccades, we divided trials into five groups: control (no light pulse), T0, T10, T20 and T30. We aligned the position, velocity and acceleration traces on saccade onset (the time at which eye velocity exceeded 15 °/s) and averaged them within each group for each saccade direction. The light pulse onset time was defined as the mean time of occurrence of the light pulse within each condition. The standard deviation of the light pulse onset time ranged from 0.54 to 1.22 ms (Tables 1 and 2).

We compared eye acceleration on control trials against those in each stimulation group using a 2-tailed Student’s t-test (Matlab “ttest2” function with default options) each millisecond from the light pulse onset time until 125 ms after saccade onset. We considered the light pulse to have had an effect on conditions in which the P values were <0.05 for at least 10 consecutive milliseconds (uncorrected for multiple comparisons). The first time at which P=0.05 was estimated using a 2-point linear interpolation. To derive an approximate confidence interval for this interpolated time, we performed a bootstrap analysis on the light-perturbed trials in each group. We obtained 1000 resampled (with replacement) data sets and compared each of these bootstrap data sets with control saccades using the procedure described above to obtain an estimate of the time of the effect. The number of samples in each bootstrap data set was equal to the number of saccades in the corresponding condition (see Tables 1 and 2).

### Normalization for population analysis

To pool data across experiments, we calculated a pair of normalization factors for each experimental session, one for rightward saccades and one for leftward saccades, that compensated for small changes in the gain of the eye coil system across days. For each saccade, we set the horizontal eye position, measured 10 ms before the identified movement onset, to zero. We then computed the mean saccade amplitude from control trials in each experiment using the eye position 125 ms after saccade onset as the endpoint. We then normalized this mean saccade amplitude of the control trials to exactly 24° (the size of the target step). We then multiplied each position, velocity and acceleration trace by the normalization factor.

### Statistics

Hypothesis testing for the comparison of saccade metrics, e.g., amplitudes, peak velocity and duration, was performed using a permutation test with 10,000 iterations (Wilcox, 2022). We considered P<0.05 as significant for all statistical tests. Averages (mean ± SD) were computed across trials within each condition. The standard deviation of the difference between the means was computed as: 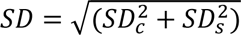 (Taylor, 1997), where *SD_c_* is the standard deviation in the control conditions and *SD_s_* is the standard deviation in the stimulation condition. Bonferroni correction was used to control the Type I error rate of multiple comparisons between control trials and the four stimulation conditions (T0, T10, T20 and T30).

### Local feedback model

We compared the effects of OMV P-cell activation with simulations from a local negative feedback model of the saccade burst generator (Fig. 13A). The model was constructed in Simulink, a graphical block diagramming tool for simulating dynamical systems in Matlab. Model simulations used a 4^th^ order Runge-Kutta ordinary differential equation solver with 0.1 ms time step. The eye plant block transfer function was (Robinson, 1964; Fuchs and Luschei, 1971; Goldstein and Robinson, 1984; Fuchs et al., 1988; Scudder, 1988):

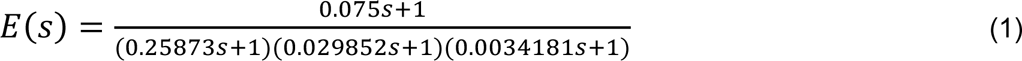

The motoneurons were implemented as a summing block that receives excitatory inputs from a position integrator (step), EBNs (pulse) and a slide component transfer function (slide) with the following weights: *wIntMN* = 0.988, *wEbnMN* = 0.104 and *wSlideMN* = 0.1092 (see Fig. 13A for weight locations in the model). The slide component transfer function was:

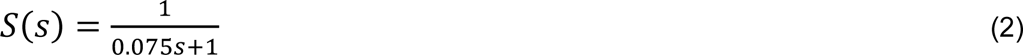

At the input of the EBN and OPN blocks, we placed a first-order low pass filter with a 2 ms time constant to simulate synaptic-input temporal integration:

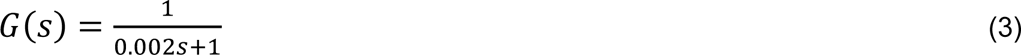

The EBN firing rate (*EBNfr*) was computed as:

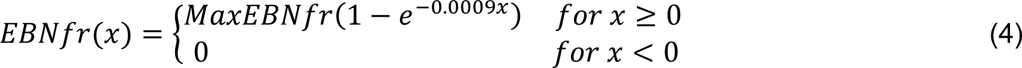

Maximum EBN firing rate (*MaxEBNfr*) was set to 1000 spikes/s to simulate fast saccades, and 750 spikes/s to simulate slow saccades. The OPN firing rate (*OPNfr*) was computed as:

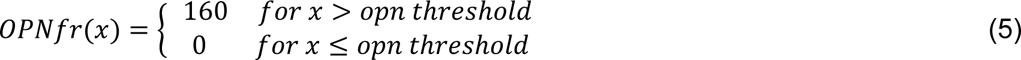

*x* in equations (4) and (5) is input firing rate.

The value of *opn threshold* was set to 50 spikes/s. The OPNs inhibited the EBNs with a weight of *wOpnEbn* = 32.

The IBNs were implemented as a summing block that receives excitatory inputs from the EBNs and the OPN trigger (150 spikes/s 10 ms pulse). The IBNs inhibited the OPNs with a weight of *wIbnOpn* = 1.50. To convert the dynamic motor error (DME) output from the comparator to a firing rate, a weight of *wCompEbn* = 80 was used. A 5-ms delay (transfer function *e*^−0.005*s*^) followed the feedback integrator 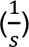 (van Gisbergen et al., 1981). All weight values were chosen to produce 24° saccades with velocity profiles that approximated data in Fig. 10.

To simulate a 24° saccade, a step was applied to the comparator, and an OPN trigger pulse occurred 40 ms later. To simulate T0 or T20 optogenetic light activation of OMV P-cells, a 450 spikes/s, 10-ms pulse occurred either 6.4 ms or 26.4 ms after saccade onset, respectively. The pulse was applied at the negative input of the EBNs, to simulate contralateral OMV stimulation, the feedback integrator, to simulate ipsilateral OMV stimulation, or both, to represent midline stimulation.

### Code accessibility

Code implementing the model is available through Github: https://github.com/robisoe/OMV_L7_paper_model

## Results

We performed 18 experiments with light pulses at four delays with respect to saccade onset (8 experiments in Sq, 10 in Tm). Significant effects on saccade metrics were obtained in all of these experiments except one (Fig. 4C, “+”) which was not analyzed further.

Recording chambers were placed at different locations relative to the functional midline in the two monkeys, and AAV injections were likely offset similarly. As a potential consequence, optical stimulation on the right side of one monkey’s OMV produced effects similar to those produced by left side stimulation in the other. For this reason, we describe saccade directions as "left" and "right" below (as opposed to the more convenient "ipsiversive" and "contraversive") to document this idiosyncrasy before reverting to the more convenient nomenclature.

### Effects of T0 light pulses on saccade metrics and kinematics

In an exemplar experiment from Sq (Fig. 3A, B), T0 light pulses did not affect the peak saccade velocity of leftward saccades (Fig. 3A, P=0.77), but caused a reacceleration ∼36 ms after pulse onset. This reacceleration prolonged the saccade by 7.18±8.41 ms, and increased saccade amplitude by 2.19±1.13°.

**Figure 3.**
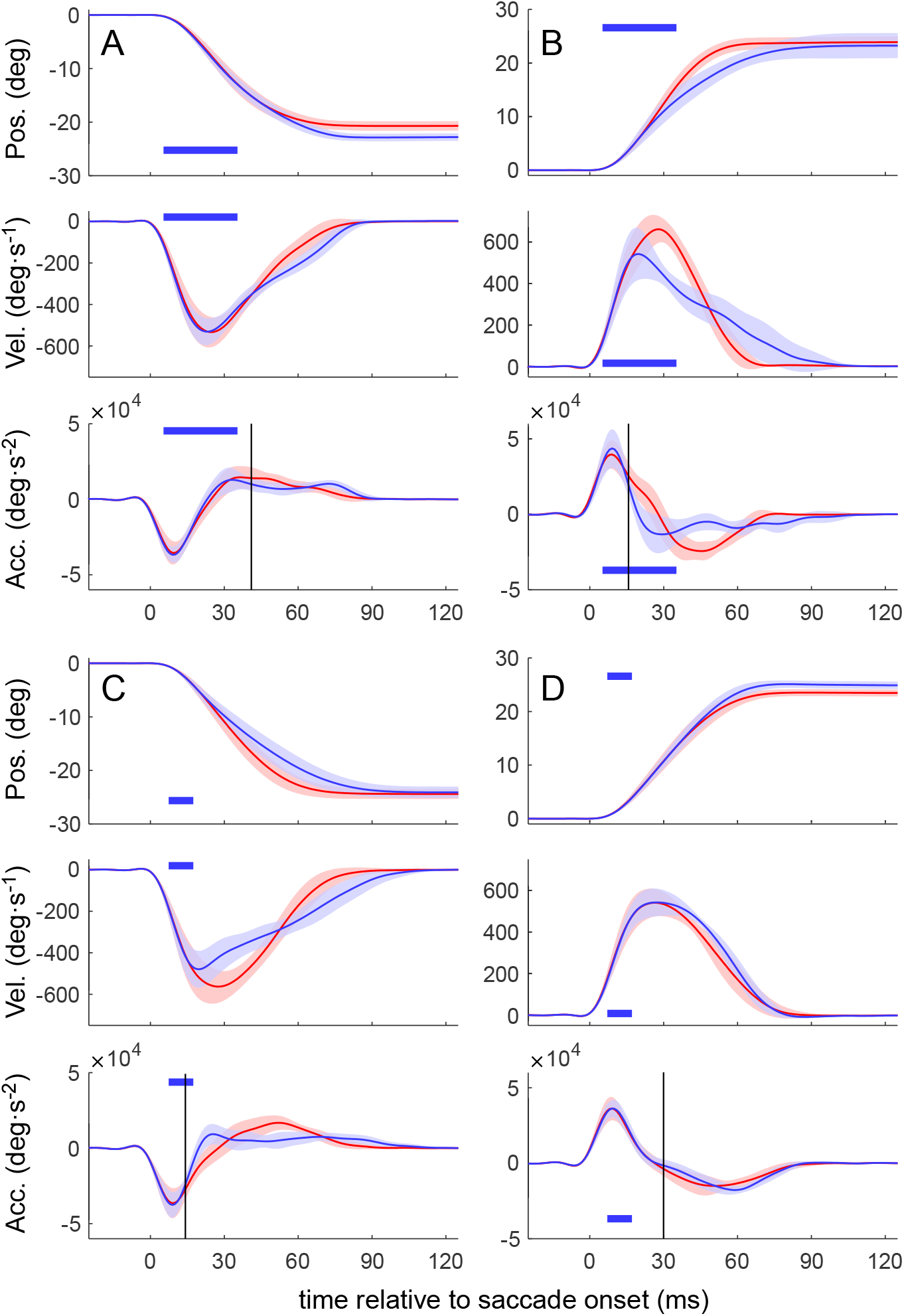
Two exemplar un-normalized data sets from animal Sq (*A* and *B)* and Tm (*C* and *D*). Top to bottom: eye position, velocity, and acceleration. Red traces, control saccades; Blue traces, saccades perturbed by T0 light pulses (blue bars); error bands = ±1 SD. Vertical lines, detected time of activation effect (time at which control and perturbed saccades differ at the P = 0.05 level). Number of trials: Sq, leftward control: 27, T0:16; rightward control: 22, T0: 22; Tm, leftward control: 34, T0: 33; rightward control: 42, T0: 27.

In contrast, T0 laser pulses (Fig. 3B) reduced the velocity of rightward saccades ∼11 ms after pulse onset, which is roughly one third of the latency on leftward saccades. Peak velocity was reduced by 120.73±127.96 °/s, (P<0.0001), but the amplitude loss during acceleration (4.49±2.36°) was compensated by an amplitude increase during deceleration (3.92±3.56°). Consequently, saccades were normometric (amplitude change relative to control: -0.57±2.68°, P=0.32).

Similar effects were obtained from animal Tm but with reversed saccade direction-dependence (Fig. 3C, D). T0 light pulses triggered by *leftward* saccades (Fig. 3C) reduced eye velocity ∼7 ms after pulse onset, decreasing peak velocity by 88.46±113.83 °/s (P<0.0001). Reduced peak velocity was followed by a reacceleration that increased the displacement during deceleration by 3.34±2.28°, which compensated the amplitude loss of 3.63±2.19° and resulted in normometric saccades (P=0.24). These data, obtained from Tm during *leftward* saccades, are similar to those obtained from Sq during *rightward* saccades (Fig. 3B).

Similarly, the effects of T0 light pulses on Tm’s *rightward* saccades (Fig. 3D) resembled those on Sq’s *leftward* saccades (Fig. 3A). Light pulses did not affect the peak velocity of Tm’s rightward saccades (P=0.92), and reacceleration started ∼23 ms after pulse onset, increasing saccade amplitude by 1.57±0.97° (P<0.0001).

As the optical fiber was moved from more central to lateral locations in the vermis of Sq, the amplitude of leftward saccades increased (Fig. 4A, left, black circles) and the amplitude of rightward saccades decreased (red circles). T0 pulses delivered near the midline (Fig. 4A, filled circles) caused both left-and rightward saccades to become slightly hypermetric (all P<0.06) by amounts that did not differ significantly (P>0.4). In Tm, as stimulation locations moved from lateral to central positions, the amplitude of rightward saccades decreased (Fig. 4A, right, red circles) and the amplitude of leftward saccades increased (black circles). At two locations just to the left of the midline (filled circles), optogenetic stimulation did not affect saccade amplitudes of either rightward or leftward saccades (P>0.1). We infer that the chamber was to the left of the functional midline in Sq and to the right in Tm. We henceforth describe Sq’s leftward saccades, and Tm’s rightward saccades, as *ipsiversive* and the opposite direction for each animal as *contraversive*. This convention is motivated by the effects of cFN inactivation: ipsiversive hypermetria and contraversive hypometria (Robinson et al., 1993; Iwamoto and Yoshida, 2002; Goffart et al., 2004; Kojima et al., 2014). Chamber locations for all experiments are shown in Fig. 4C (gray circles). For the population analyses, we pooled data from experiments conducted at the leftmost locations (Fig. 4B, left, open circles) in Sq and, separately, those conducted at the rightmost locations in Tm (Fig. 4B, right, open circles).

**Figure 4.**
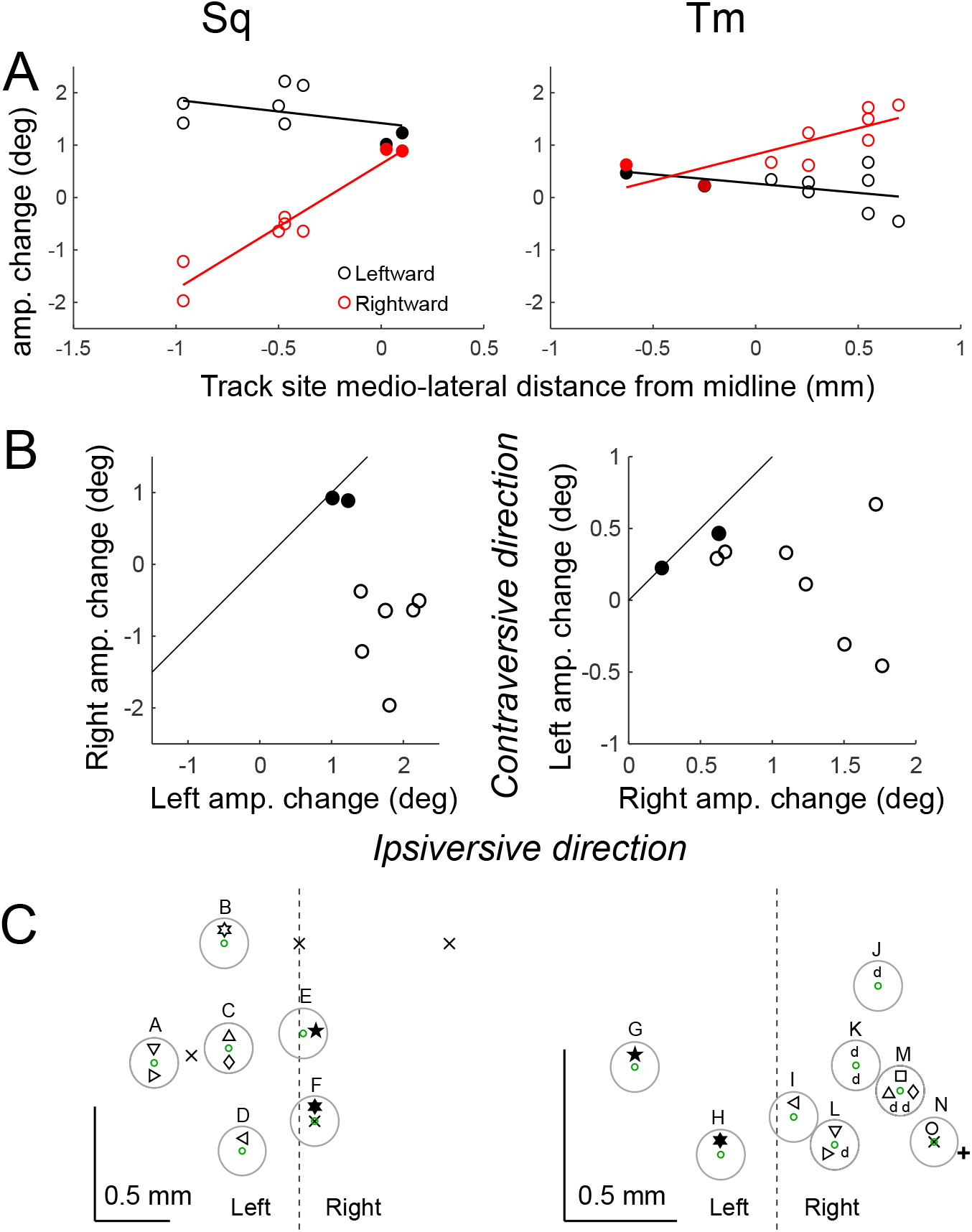
The effects of optical fiber location on saccade amplitude change. Sq, left column; Tm, right column. *A*, Saccade amplitude change vs. medio-lateral location of the optical fiber in the recording chamber. Red circles, rightward saccades; black circles, leftward saccades. *B*, amplitude change of rightward and leftward saccades. Filled circles, fiber located to the right (for Sq) or to the left (for Tm) of midline. *C*, Actual locations of the optical fiber for all experiments (green circles); AAV injection sites (X). Other symbols represent individual experiments. Data in Figs. 3A and 3B were collected from recording site D. Data in Figs. 3C and 3D were collected from recording site M. The “d”s represent 6 experiments using variable light pulse durations. +, an experiment that produced no significant change in saccade either saccade metrics or kinematics.

### Effects of laser pulse time on ipsiversive saccades

Optical stimulation affected the deceleration phase of ipsiversive saccades, largely irrespective of pulse timing. Despite the 10 ms delay between them, T0 and T10 pulses produced reaccelerations that occurred <2 ms apart (Fig. 5A, C, red and green vertical lines; Table 1; P= 0.72 and 0.031 for Sq and Tm, respectively, permutation test of onset comparison, 10,000 iterations). T20 and T30 pulses affected saccades slightly later, but still within 20 ms of the perturbations produced by T0 pulses (Fig. 5B, D, blue and orange vertical lines; Table 1; P <0.041 for comparison across T0 to T20, and P <0.002 across T0 to T30, permutation test of onset-slope comparison, 10,000 iterations). In summary, optical stimulation of the ipsilateral OMV produced a reacceleration that was more closely time-locked to the movements than it was to the timing of the light pulses. This result suggests that the access of the ipsilateral OMV to oculomotor premotor burst neurons is indirect, affecting saccades only after a relatively long delay that depends on the state of the movement.

**Figure 5.**
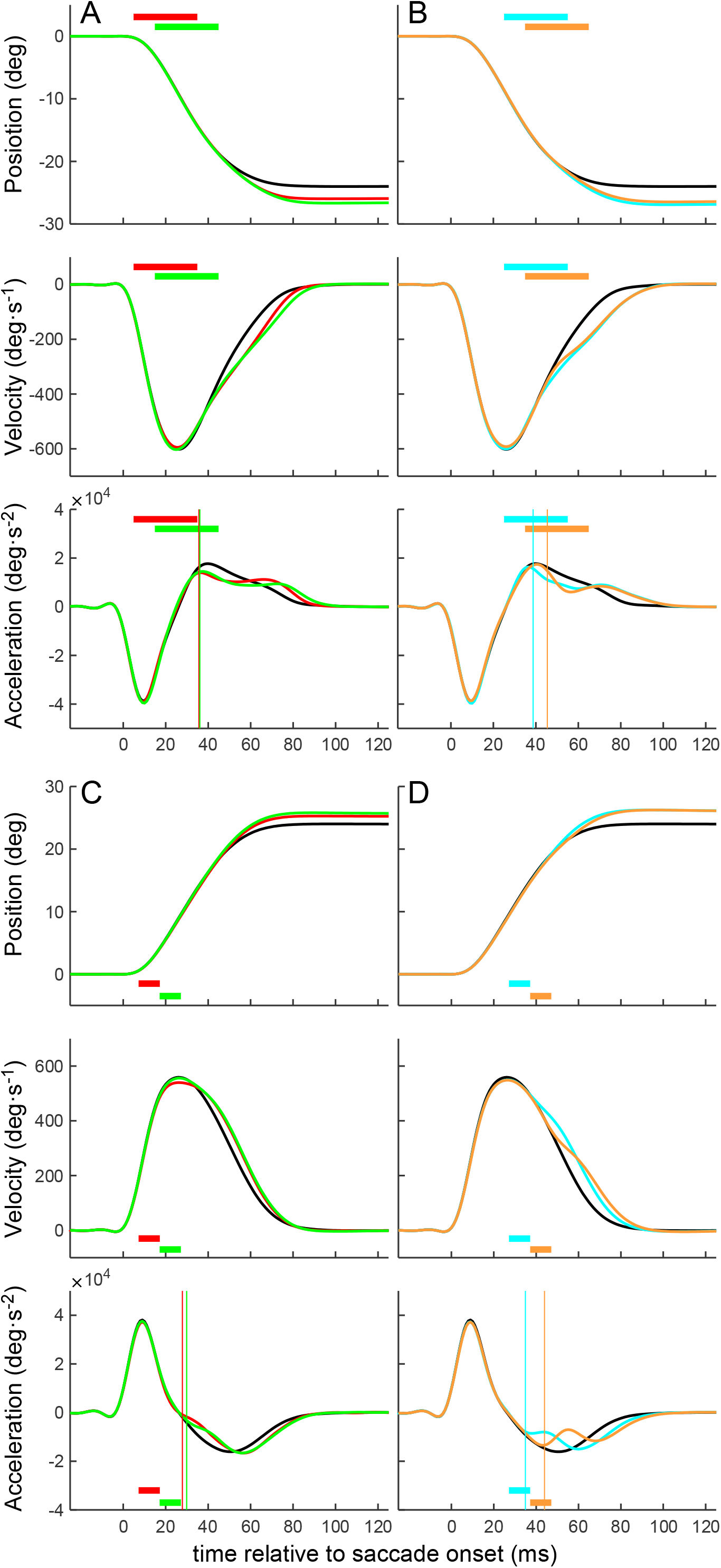
Population averages of *ipsiversive* saccade time courses. Traces are normalized (see Materials and Methods). *A* and *B*, data from animal Sq. Top to bottom, eye position, velocity and acceleration. Horizontal bars, light pulses. Black, control saccades; red, T0 pulse perturbed saccades; green, T10 pulse perturbed saccades; blue, T20 pulse perturbed saccades; orange, T30 pulse perturbed saccades. Solid vertical lines indicate the time of P=0.05 difference from the control. *C* and *D*, data from animal Tm using the same conventions as in *A* and *B*.

### Change in saccade metrics

Optical stimulation caused ipsiversive saccades to become hypermetric (Fig. 5A-D, top panels; Table 1; P<0.0001, both animals), but did not affect peak velocity (one-way ANOVA: F<1.5, P>0.1, both animals), acceleration duration (F<1.7, P>0.1, both animals), or displacement amplitude of the eye during the acceleration phase (F<1.3, P>0.1, both animals). It did, however, increase deceleration duration (P<0.0001, all conditions except for T0 and T10 in Tm for which P>0.15).

### Titrating the reacceleration effects

To test the hypothesis that the ipsilateral OMV/cFN complex is part of the integrator feedback pathway, we asked whether longer pulses increased saccade amplitudes. A positive result would suggest that the perturbed activity is integrated. We fixed the start time of the laser pulse at T0 and varied its duration from 1.25 to 10 ms (experiments conducted in Tm only). Despite their brevity, 1.25-ms pulses extended ipsiversive saccades by 0.75±0.92° (P<0.0001), demonstrating the remarkable effectiveness of this optogenetic manipulation on primate behavior (Fig. 6A, B). Consistent with this hypothesis, increasing the light pulse duration produced yet larger saccades (Fig. 6B, red line, slope: 0.22 °/log_2_[pulse duration], P<0.0001).

**Figure 6.**
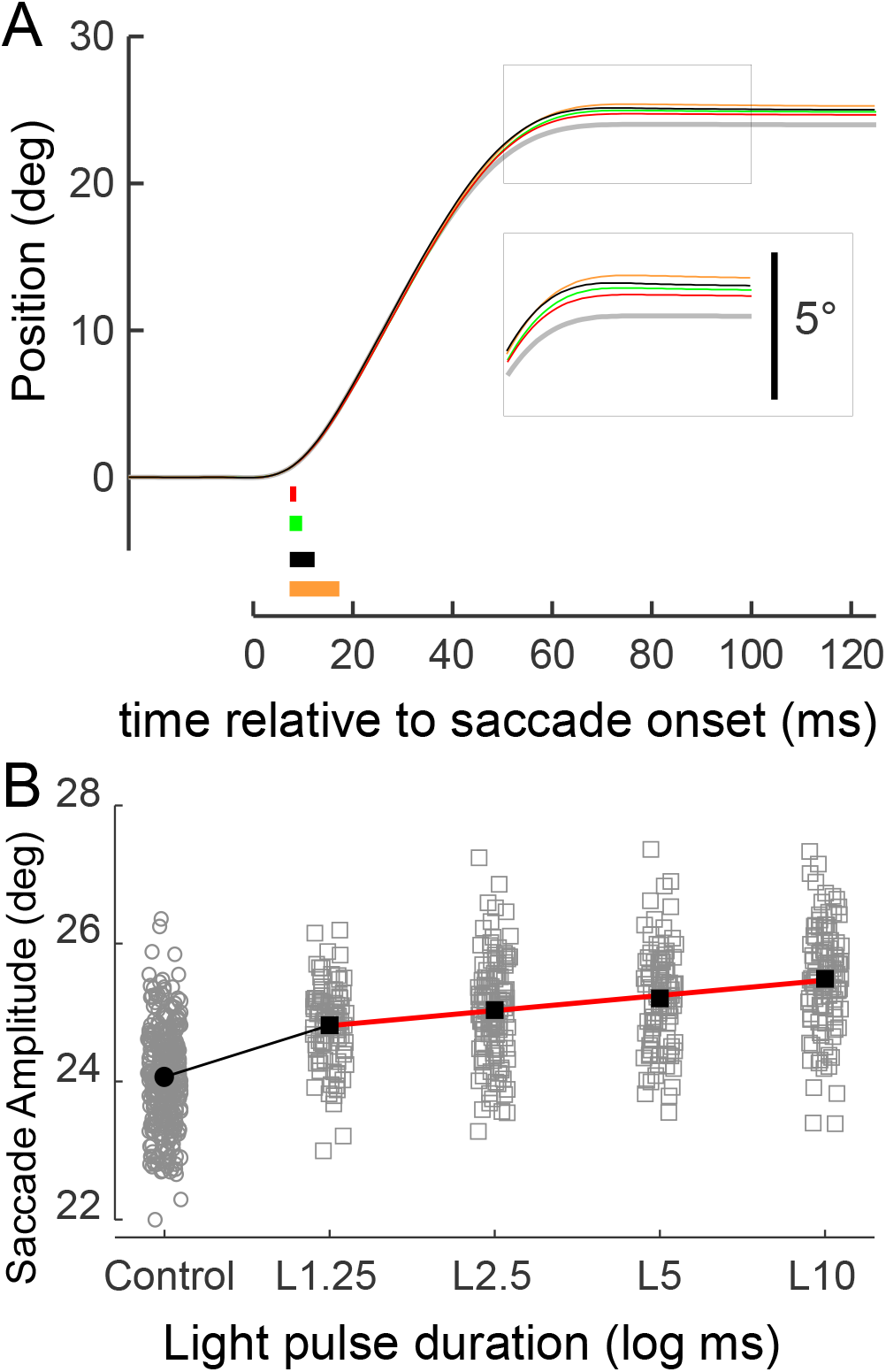
Population average effects of light pulse duration on ipsiversive saccade amplitudes. *A*, time courses of saccades after variable duration pulses starting at T0. Horizontal bars, light pulses; pulse durations: red, 1.25 ms; green, 2.5 ms; black, 5 ms; orange, 10 ms. *B*, saccade amplitude vs. pulse durations (squares). Circles, control. Filled circle and squares, averages. Red lines, linear regression. N=328, 93, 120, 98 and 109 for control and pulse durations of 1.25, 2.5, 5 and 10 ms, respectively. Points were jittered to facilitate visualization.

In summary, brief activation of ipsilateral OMV P-cells reaccelerated saccades during the deceleration phase and produced hypermetria. The time at which reacceleration occurred depended on the state of the movement but relatively little on stimulation timing. Hypermetria increased with stimulation duration, consistent with an integration mechanism either at or downstream of the P-cells. The integration process starts early, around the time of saccade onset, and its effect appears >20 ms later. These results contrast with the effects of OMV P-cell activation on contralateral saccades, which we describe below.

### Effects of laser pulse time on contraversive saccades

The main effect of the light pulse on contraversive saccades was a deceleration which was followed by a compensatory reacceleration (Fig. 3B, C). The timing of this deceleration was strongly dependent on the timing of the light pulse. For Sq, T0, T10, and T20 pulses decelerated saccades at short, relatively fixed latency with respect to laser onset (∼6–9 ms) and consequently at long, variable latency with respect to saccade initiation (Fig. 7A, B; Table 2). Similar results were obtained for Tm; in this animal, T0, T10, T20, and T30 pulses decelerated saccades ∼4–5 ms after laser onset (Table 2; Fig. 7C, D, vertical lines). These results show that OMV stimulation decelerates contraversive saccades at low latency irrespective of the state of the movement. This low latency contrasts with the longer latency effects produced by ipsilateral stimulation and suggests that contralateral P-cells have more direct access to the SBG premotor burst neurons than the ipsilateral P-cells do.

**Figure 7.**
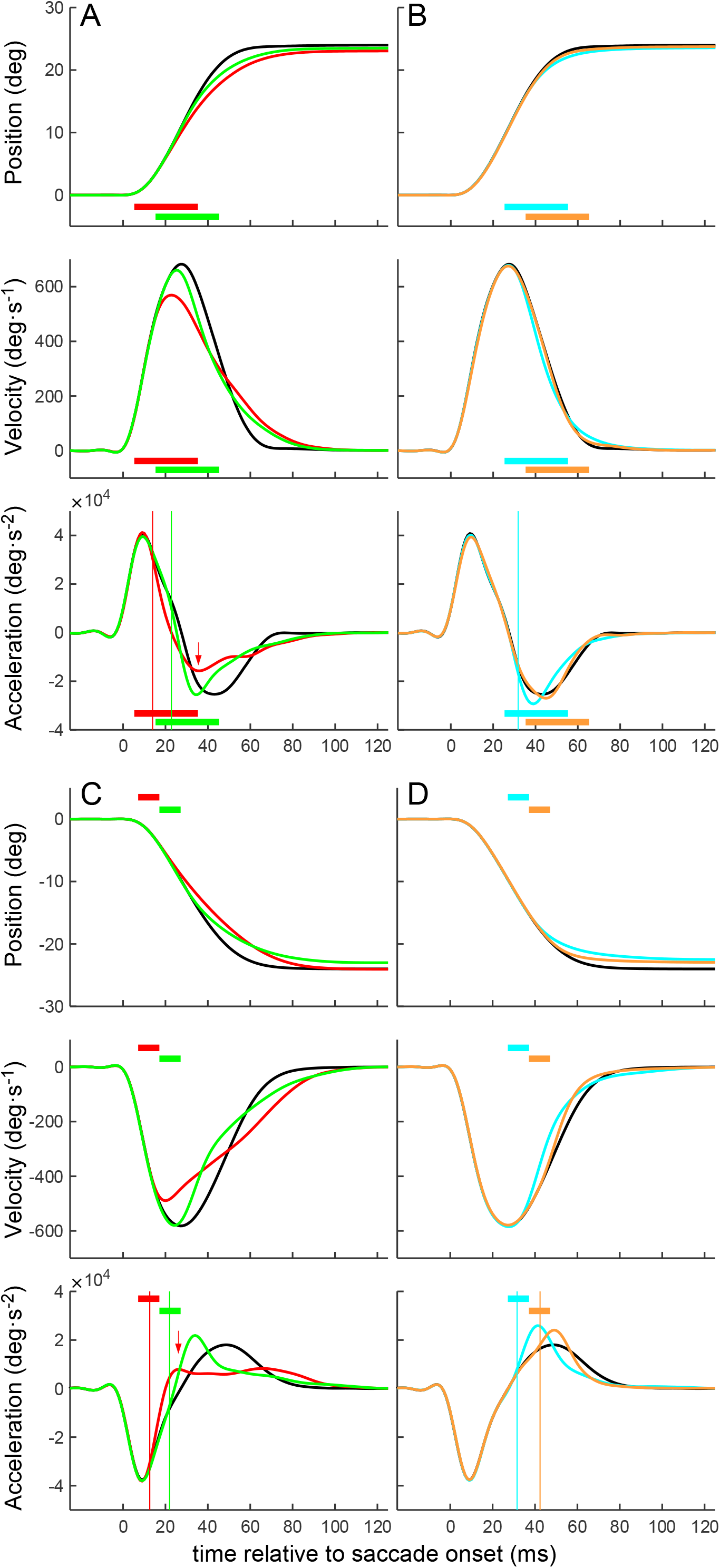
Population average contraversive saccade time courses. Conventions are as in Fig. 5. *A* and *B*, data from animal Sq. *C* and *D*, data from animal Tm.

### Change in Saccade Metrics

T0 pulses reduced the peak velocity of contraversive saccades by 106.61±130.27 °/s in Sq and by 97.43±114.27 °/s in Tm, (both P<0.0001; Fig. 7A, C, velocity panels, red traces; Table 2). Under most circumstances, lower velocities are associated with smaller saccades—a relationship known as the main sequence (Bahill et al., 1975). This relationship predicts that a 10% decrease in the peak velocity of a 24° saccade should produce a 5° reduction in saccade amplitude (Fuchs, 1967). To the contrary, the amplitudes of contraversive saccades remained relatively unchanged despite the nearly 15% reduction of peak velocity produced by optogenetic OMV P-cell stimulation. We note that saccade amplitude decreased by 0.93±2.18° in Sq (P<0.001, Fig. 7A, position panel, red trace), but this decrease is small relative to that expected from the main sequence. No significant change was found in Tm (P=0.39, Fig. 7C, position panel, red trace; Table 2).

T0 optogenetic stimulation abbreviated the acceleration phase of contraversive saccades, decreasing eye displacement during this phase by 3.07±2.51 and 3.77±2.17° in Sq and Tm, respectively (Table 2). The eyes reaccelerated during the deceleration phase (35.83±0.95 and 26.18±0.39 ms after saccade onset for Sq and Tm, respectively; Fig. 7A and C, acceleration panels, red traces) and therefore landed more or less on target. Similar results were obtained from experiments in which the duration of the T0 light pulse was varied from 1.25 to 10 ms (Tm only). Longer light pulses more profoundly reduced the eye displacement during the acceleration phase of the saccade, but this was compensated by yet longer deceleration phases so that, in all cases, saccade amplitude was nearly constant (one-way ANOVA across control and all 4 pulse duration groups: F(4, 803)=2.08, P=0.081). These data attest to the engagement of a mechanism that extends the deceleration phase to compensate for an abbreviated acceleration phase.

The negative feedback model (Fig. 1B) predicts that, for a given saccade amplitude, saccade duration should decrease as peak velocity increases (Soetedjo et al., 2002; Barton et al., 2003). Linear regression confirmed this prediction (Fig. 8A–D, black circles and lines). Light pulses applied at T0 or T10 maintained the linear relationship between saccade duration and peak velocity but steepened it (Fig. 8A–D, red and green lines). These effects are consistent with the negative feedback model.

**Figure 8.**
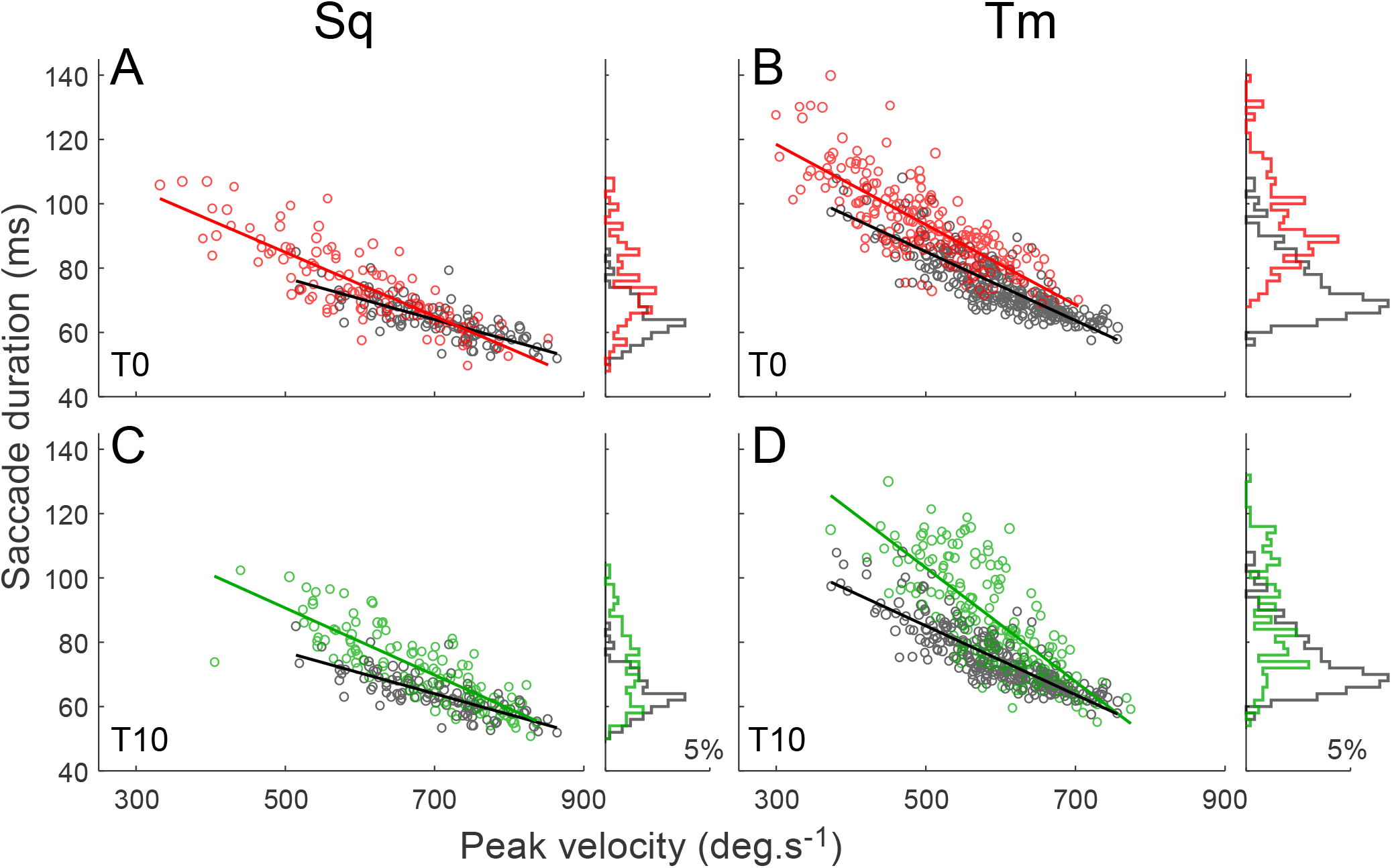
Duration vs. peak velocity of contraversive saccades following T0 and T10 pulses. *A – D*, T0 (red) and T10 (green) perturbed saccades from Sq (A, C) and Tm (B, D); control saccades (black). Linear fits: black lines: R^2^=0.61 and 0.73, for Sq and Tm, respectively; red lines: R^2^=0.70 and 0.62, for Sq and Tm, respectively; green lines: R^2^=0.69 and 0.58, for Sq and Tm, respectively. Histogram of saccade durations (right column).

On T20 and T30 trials, slow saccades had unusually long deceleration durations, and the relationship between peak velocity and deceleration duration on stimulated trials was better described by a second-order polynomial than a line (Fig. 9A-C, blue and orange data and fits, F-test, P<0.0001). Deceleration duration decreased steeply with increasing peak velocity and was even shorter on some stimulation trials than on control trials. These results suggest that late OMV P-cell activation truncated the fastest contraversive saccades and prolonged the deceleration phase of the slower ones. Indeed, an analysis of saccade velocity profiles showed that T20 and T30 pulses decelerated contraversive saccades, and truncated the fast ones selectively (Fig. 10). An analysis of saccade amplitudes showed that fast saccades became hypometric (Fig. 10A-C, insets, all P values <0.01) whereas slow saccades remained near normometric (Fig. 10D-F, insets, all P values >0.2). These results suggest that the effects of late contralateral OMV stimulation depend on the state of the movement. We propose a mechanism based on dynamic motor error for this dependence in *Comparison with model simulations*.

**Figure 9.**
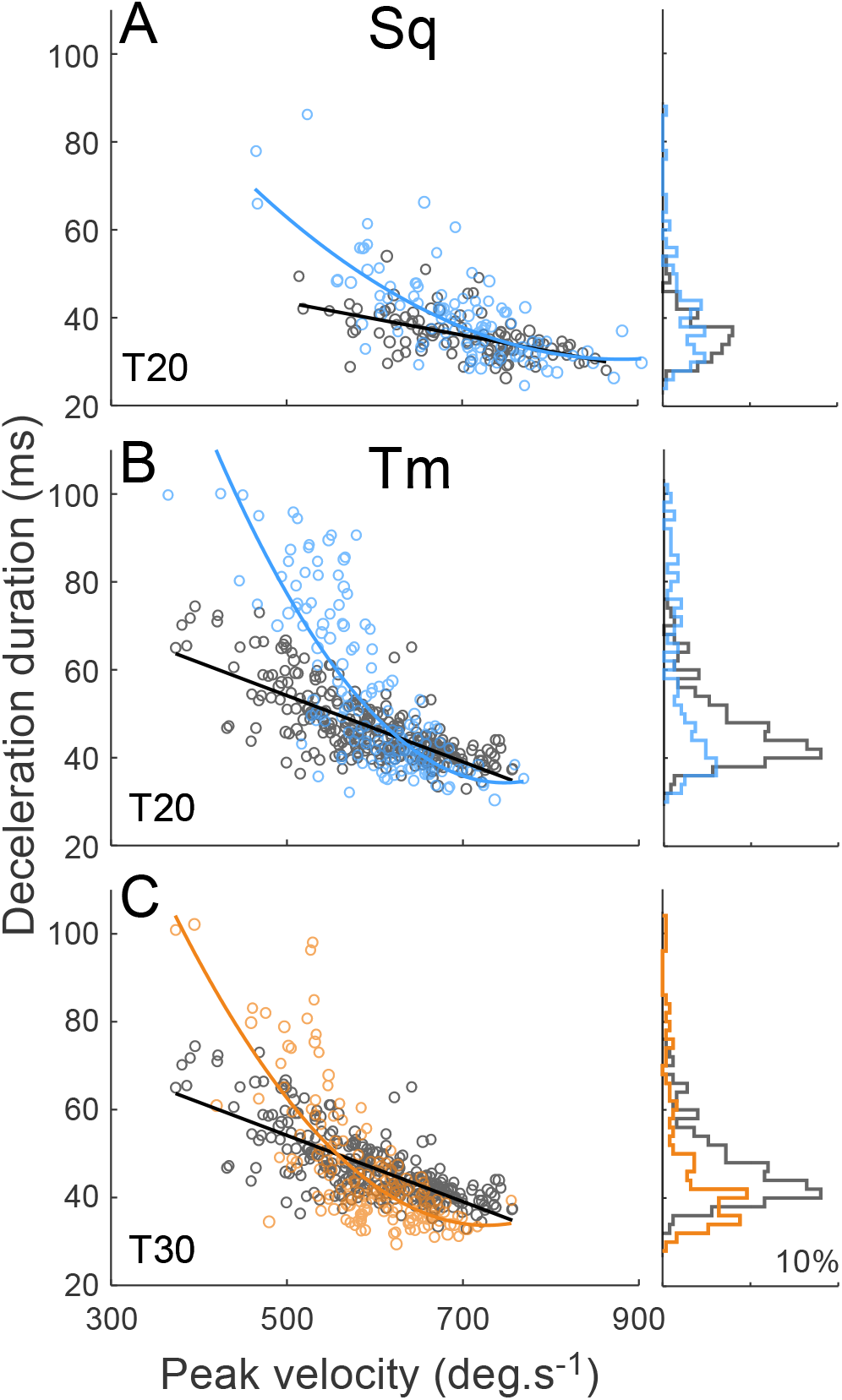
Deceleration duration vs peak velocity of contraversive saccades following T20 and T30 pulses. *A – C*, T20 (blue) and T30 (orange) perturbed saccades from Sq (A) and Tm (B, C); control saccades (black). Black lines are linear fits; curved blue and orange lines are second-order polynomial fits. Histogram of saccade deceleration durations (right column).

**Figure 10.**
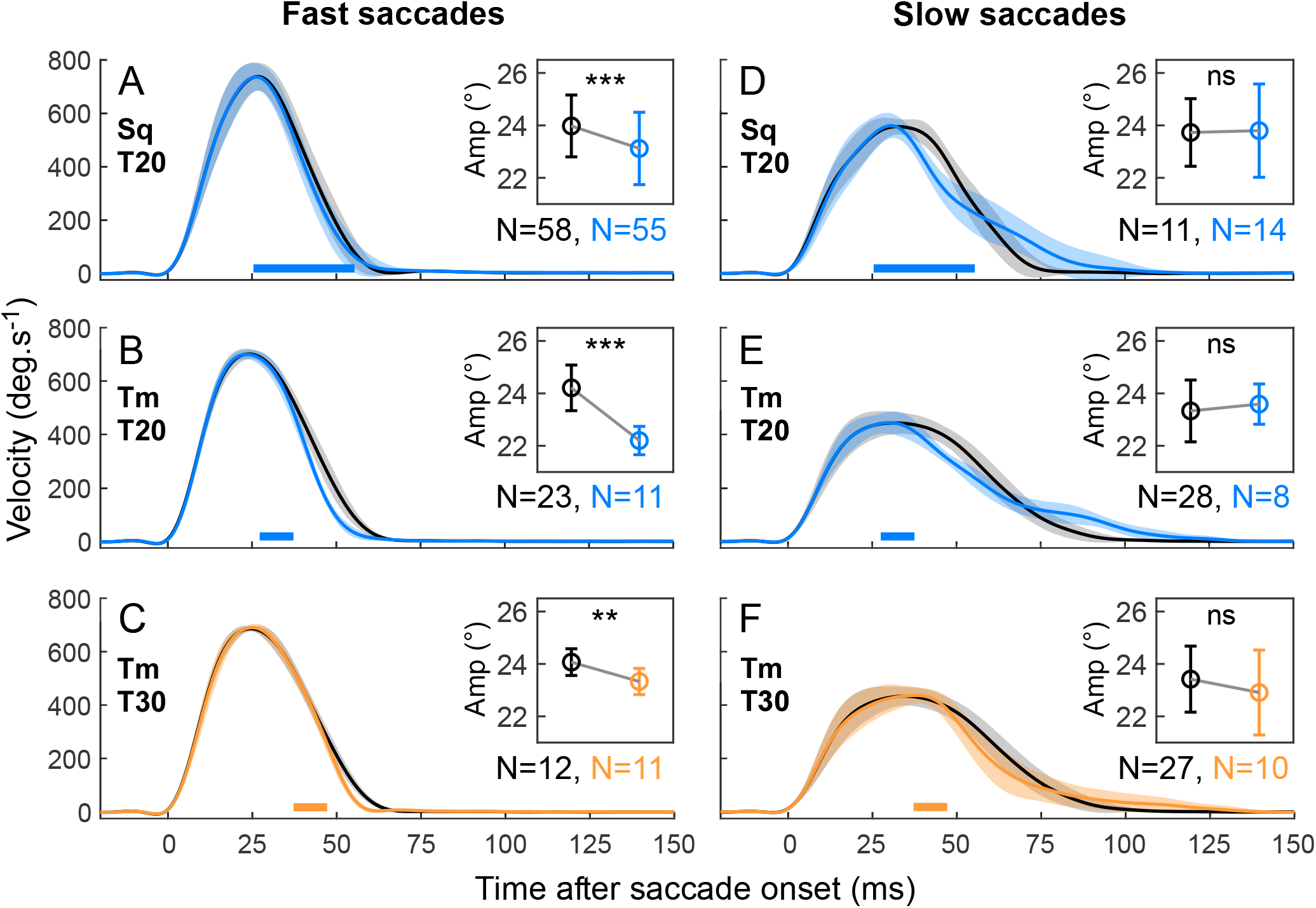
Comparison of effects of T20 and T30 pulses on fast (*A – C*) and slow (*D – F*) contraversive saccades. *A* and *D*, monkey Sq, T20 data; fast saccades (A) 700*–*900 °/s; slow saccades (D) <600 °/s. *B* and *E*, monkey Tm, T20 data; fast saccades (B) 680*–*750 °/s; slow saccades (E) <500 °/s. *C* and *F*, monkey Tm, T30 data; fast saccades (C) 680*–*760 °/s; slow saccades (F) <496 °/s. Black, control; blue and orange, T20 and T30 data, respectively. Error band = ±1 SD. Insets, saccade amplitude comparisons. Error bar in inset = ±1 SD. Velocities are plotted in the positive direction for both animals. Trials were selected to match the time course of saccade acceleration phase.

### Mixed contra-and ipsiversive effects after midline stimulation

In four experiments (two on Sq and two on Tm), we delivered light pulses near the functional midline of vermis (Fig. 4A, B, filled circles). In presenting these effects, we maintain the convention that "ipsiversive" is left for Sq and right for Tm.

In both animals, T0 light pulses to the midline reduced saccade peak velocity (Fig. 11A, C, E and G, middle panels, red traces, P<0.0001), which is consistent with the contraversive effect. This was followed by reacceleration that produced small, but significant, hypermetria, consistent with the ipsiversive effect (1.23±1.54, 0.41±1.47, 1.08±1.99 and 0.38±1.35°; Fig. 11A, C, E and G, respectively, P<0.031). T10 pulses also decelerated saccades (green traces) at short latency and caused a reacceleration at longer latency that produced hypermetria (Fig. 11A, C and G; P<0.0001). Similar results were produced by T20 and T30 stimulation pulses (Fig. 11B, D, F and H). Hypometria was not produced by any pulse condition in either animal. The mixed contra-and ipsiversive effects are consistent with the simultaneous activation of P-cells on both sides of the OMV, an interpretation supported by analysis in *Comparison with model simulations*.

**Figure 11.**
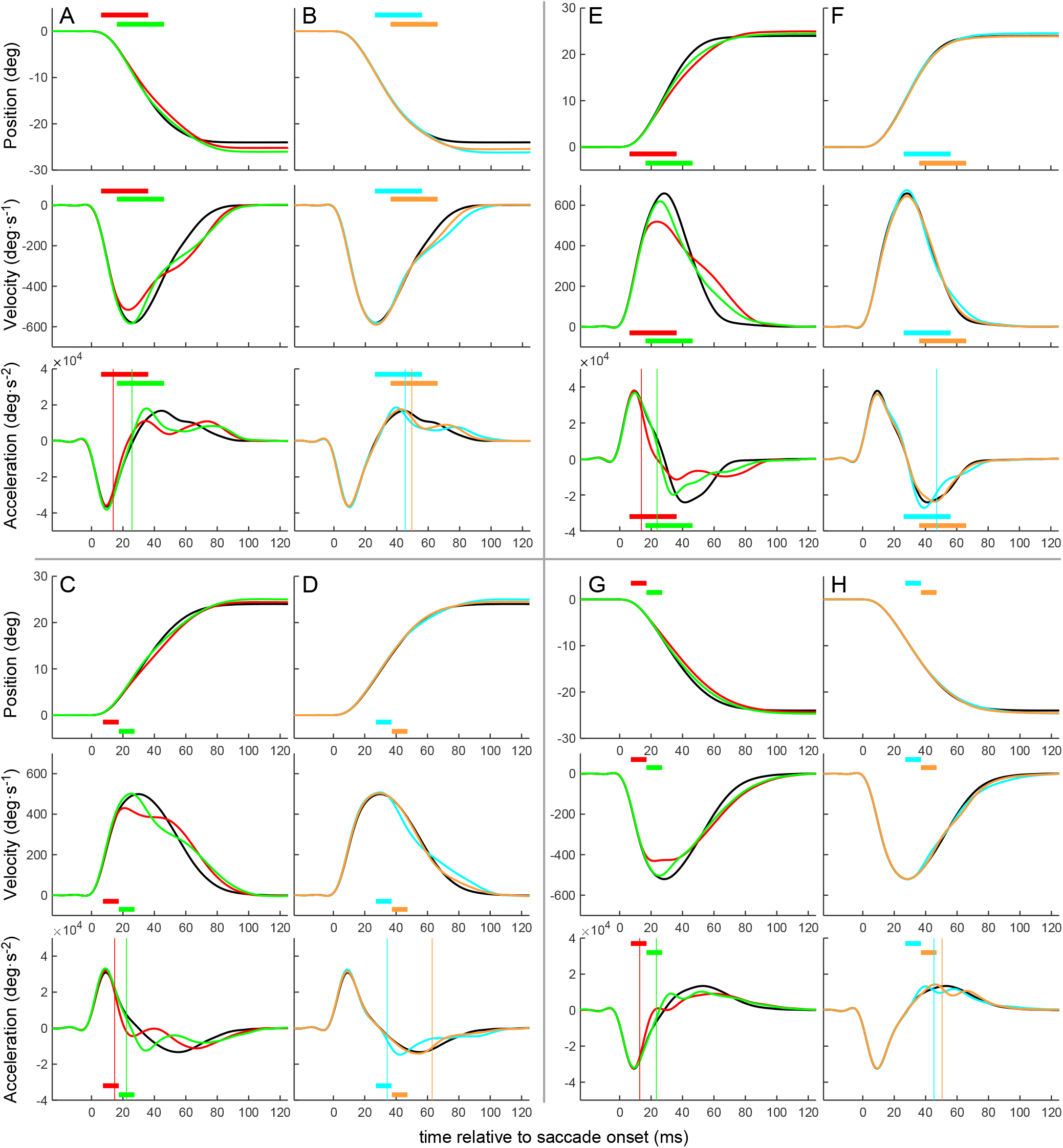
Population average saccade time courses following midline OMV stimulation. Conventions are as in Figs. 5 and 7. *A* and *B*, leftward saccades from animal Sq, N=50, 39, 46, 38, and 36 for control and T0, T10, T20 and T30, respectively. *C* and *D*, rightward saccades from animal Tm, N=84, 48, 43, 26, and 45 for control, and T0, T10, T20 and T30, respectively. *E* and *F*, rightward saccades from animal Sq, N=35, 34, 34, 29, and 26 for control, and T0, T10, T20 and T30, respectively. *G* and *H*, leftward saccades from animal Tm, N=123, 87, 60, 78, and 66 for control, and T0, T10, T20 and T30, respectively.

### Does the ipsilateral OMV help compensate for the effects of contralateral activation?

T0- perturbed contraversive saccades exhibited a compensatory reacceleration during the deceleration phase that brought the saccades close to normometric (Fig. 7; Table 2). An intriguing possibility is that the ipsilateral OMV/cFN was responsible for the compensatory reacceleration of perturbed contraversive saccades.

To test this possibility, we compared the time of occurrence of the ipsiversive effects (Fig. 12A, B, respectively) with the time of the compensatory reacceleration of perturbed contraversive saccades (Fig. 12C, D, respectively). We analyzed T0 stimulation trials because they exhibited the earliest and largest changes in velocity.

**Figure 12.**
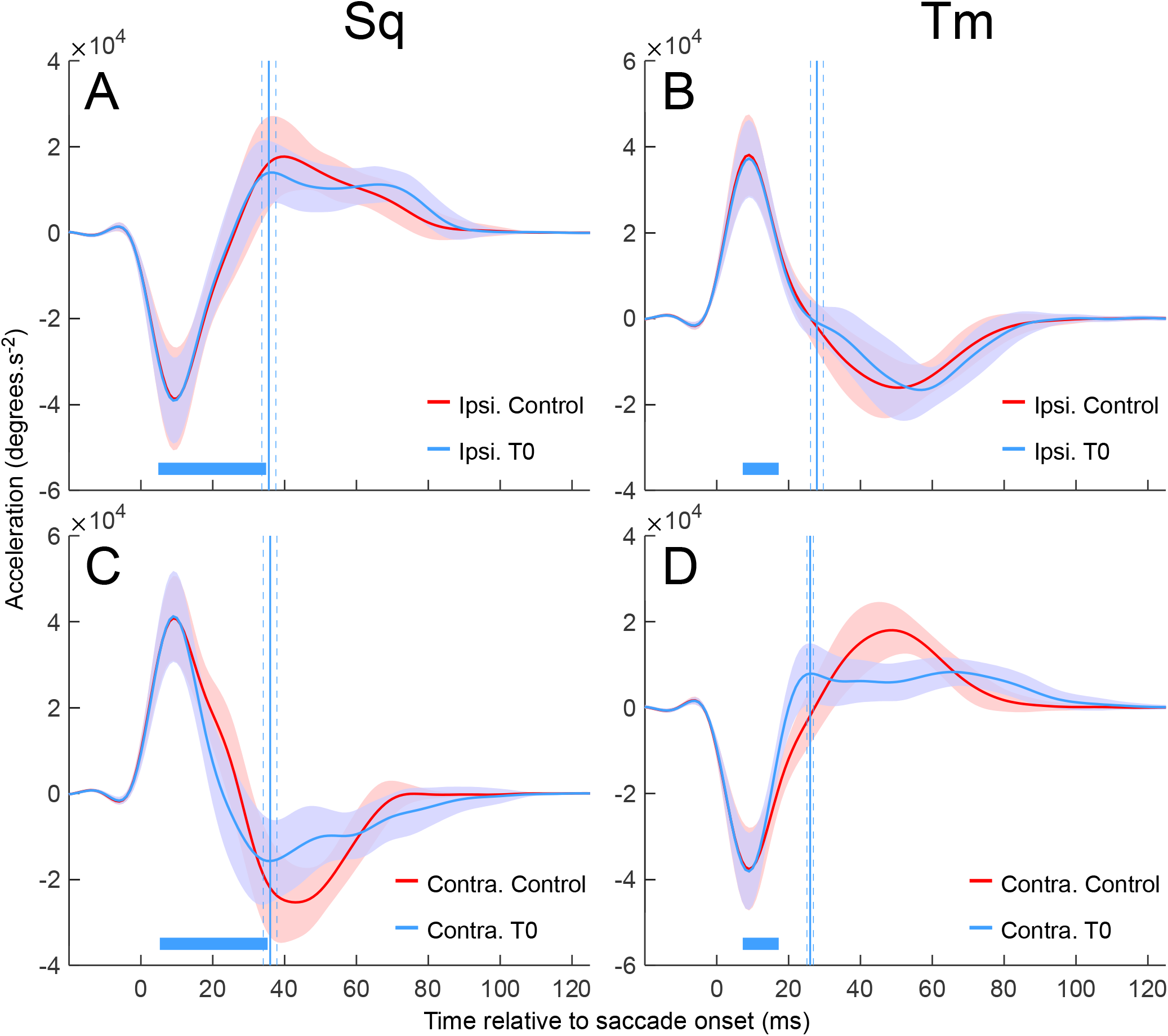
Timing comparison between T0 ipsiversive reacceleration and T0 contraversive compensatory reacceleration. *A* and *B*, ipsiversive acceleration time courses of Sq and Tm, respectively. Red, control ipsiversive saccades; blue, T0 perturbed ipsiversive saccades. Error bands = ±1 SD. Solid vertical lines, mean time of occurrence of the ipsiversive reacceleration effect, dashed lines, ±2 SE. Red traces in A and B are identical to black traces in the bottom panel of Fig. 5A and 5C, respectively. Blue traces in A and B are identical to red traces in the bottom panel of Fig. 5A and 5C, respectively. *C* and *D*, contraversive acceleration time courses of Sq and Tm, respectively. Red, control contraversive saccades; blue, T0 perturbed contraversive saccades. Solid vertical lines, mean time occurrence of contraversive compensatory reacceleration. Red traces in C and D are identical to black traces in the bottom panel of Fig. 7A and 7C, respectively. Blue traces in C and D are identical to red traces in the bottom panel of Fig. 7A and 7C, respectively.

The compensatory reacceleration in Sq occurred 35.83±0.95 (mean±SE) ms after the onset of perturbed contraversive saccades (Fig. 12C, blue line). For comparison, the effect of T0 pulses on ipsiversive saccades occurred 35.62±0.97 ms after saccade onset (Fig. 12A, blue line). These two times differ by <1 ms. The same comparison for Tm yielded similar results; compensatory reacceleration occurred 26.18±0.39 ms after the onset of perturbed contraversive saccades (Fig. 12D, blue line), and T0 pulses affected ipsiversive saccades 27.89±0.89 ms after saccade onset (Fig. 12B, blue line). These results are consistent with a role for the ipsilateral OMV/cFN in the circuit that compensates for the decrease in the velocity produced by contralateral OMV P-cell activity.

**Figure 13.**
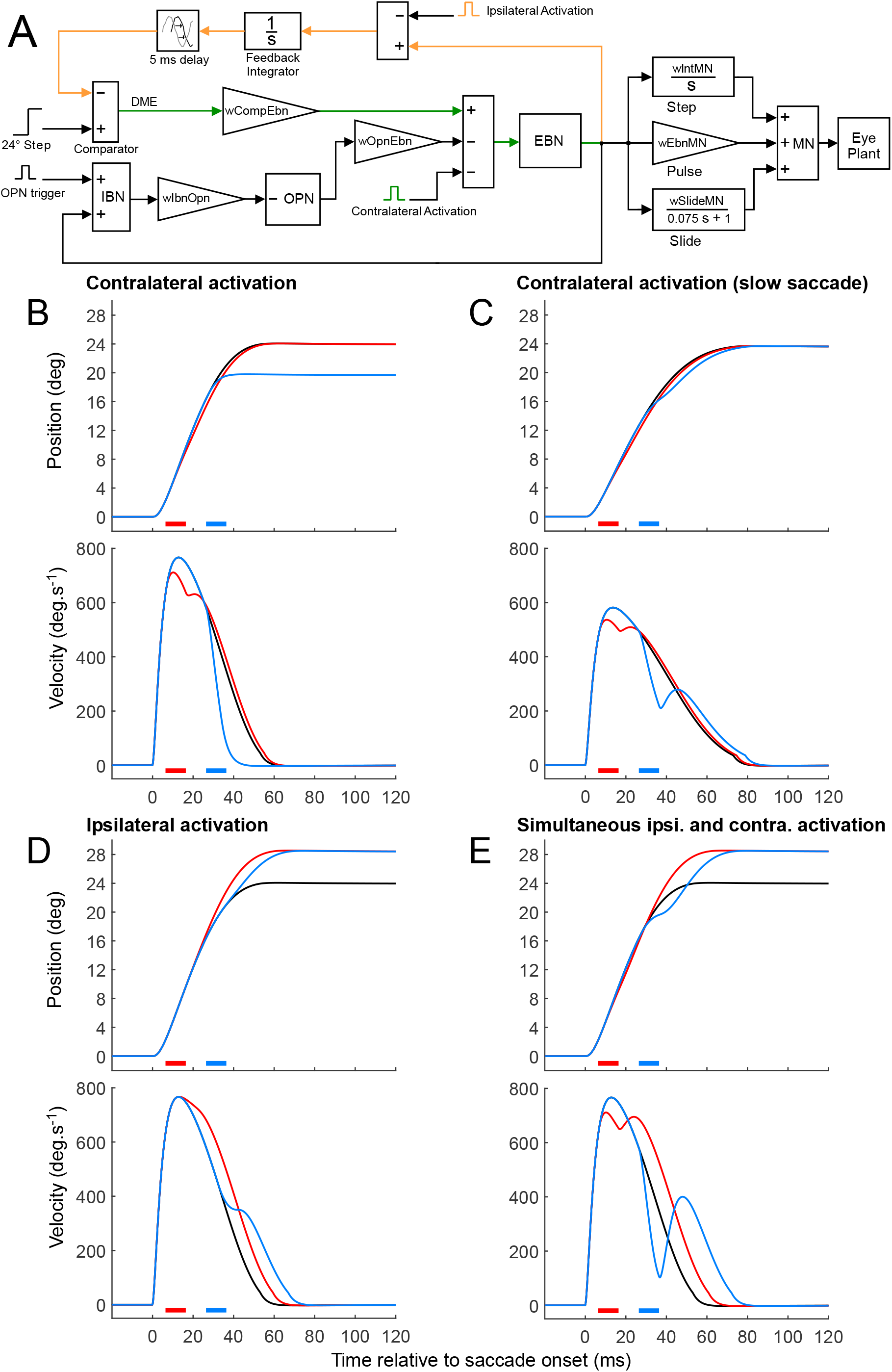
Model and simulations. *A*, The local feedback model of the saccade generator. Abbreviations, DME, dynamic motor error; IBN, inhibitory burst neurons; OPN, omnipause neurons; EBN, excitatory burst neurons; MN, motoneurons. See Material and Methods for equations and weight values. Green pulse, contralateral P-cell activation, which inhibits the EBNs. Orange pulse, ipsilateral P-cell activation, which inhibits the feedback integrator. *B*, Simulations of contralateral activation, top panel: position, lower panel, velocity. Black, control; red, early activation pulse; blue, later activation pulse. Bars, activation pulses. *C*, the same as in B, but for slower saccades (*MaxEBNfr* = 750 spikes/s, see Materials and Methods). *D*, Simulations of ipsilateral activation. E, Simulations of simultaneous ipsi-and contralateral activation.

### Local feedback model

The distinct effects of OMV P-cell stimulation on contra-and ipsiversive saccades suggest that the two sides of the OMV play distinct roles in the local feedback model of saccade production (Fig. 1B). The short latency deceleration and compensatory reacceleration produced by contralateral OMV stimulation are consistent with a role in the feedforward pathway. The scaling of saccade amplitude with the duration of ipsilateral OMV stimulation and the long latency of these effects suggest a role in the feedback pathway. Further evidence for these hypothesized roles is the similarity in timing between the contraversive compensatory reacceleration and the stimulation-induced ipsiversive reacceleration.

### Comparison with model simulations

We formalized these intuitions using a local feedback model of the saccade generator that included a “latch” circuit to prevent the OPN from firing during a saccade (Fuchs et al., 1985; Takahashi et al., 2022; Fig. 13A, see Materials and Methods for detail quantitative description of the model). The input to the model is a 24° displacement command, which is compared with the estimate of eye displacement provided by the feedback integrator to produce a dynamic motor error (DME). The saccade continues until the DME reaches zero.

During fixation, the OPNs inhibit both EBNs and inhibitory burst neurons (IBNs) (Langer and Kaneko, 1983; Iwamoto et al., 2009). A saccade begins when a trigger signal (10 ms black pulse, 150 spikes/s), presumably from the superior colliculus (Sparks et al., 2000), inhibits the OPNs via the IBNs (Takahashi et al., 2022) and allows the EBNs to respond to the DME. The EBNs drive the abducens motoneurons (MNs) directly to produce the burst that accelerates the eyes. To prevent post-saccadic drift, the signal carried by the EBNs is integrated (“Step” term) and low pass filtered (“Slide” term) before reaching the MNs (Collins et al., 1975; Optican and Miles, 1985; Kaneko, 1997; Sylvestre and Cullen, 1999). The EBNs also drive the IBNs (Sasaki and Shimazu, 1981; Strassman et al., 1986) which in turn, inhibit the OPNs. The EBN-IBN-OPN-EBN loop forms a latching circuit that maintains the OPN’s pause during a saccade.

The cFN, which projects to both the contralateral EBNs and IBNs, is inhibited by OMV P-cells. Consistent with previous local feedback models in which the IBNs are part of the OPN latching circuit (van Gisbergen et al., 1981; Scudder, 1988), optical activation of the contralateral vermis was implemented as a 10 ms, 450 spikes/s pulse that inhibited the EBNs only. How the ipsilateral cFN influences the burst generator is less clear because its targets, the contralateral EBNs and IBNs, are silent during ipsiversive saccades (Strassman et al., 1986; Scudder et al., 1988; Noda et al., 1990). The effect of ipsilateral OMV P-cell stimulation in our data appeared to be integrated and produced hypermetria, so we modeled it as a 10 ms, 450 spikes/s pulse (orange) with a negative weight to the feedback integrator.

Under this model, activating contralateral OMV P-cells (Fig. 13B, red bar) produces a deceleration at short latency (Fig. 13B, velocity panel, compare black control and red perturbed saccades). Shortly after the end of this pulse, the saccade reaccelerates and lands on target due to the feedback circuit (Fig. 13B, position and velocity panels, red trace). Similar effects occur for fast and slow saccades (Fig. 13C, red traces).

Contralateral activation during the deceleration phase (26.4 ms after saccade onset, Fig. 13B, C, blue bar) affect simulated fast and slow saccades in different ways. Both types of saccades are decelerated by the pulse, but slow saccades reaccelerate and compensate fully for the deceleration (Fig. 13C, blue trace). Fast saccades do not reaccelerate and consequently become hypometric (Fig. 13B, blue trace).

The reason for this difference is that the DME decreases quickly during a fast saccade and slowly during a slow saccade. The slower decrease of the DME slows the decrease of EBN activity, which prevents the OPN input from exceeding the 50 spikes/s threshold. Consequently, the OPNs remain paused and allow the eye to continue to move. Fast saccades, in contrast, have low DME, and therefore low EBN activity, at the time of the pulse. The additional suppression of EBNs afforded by the pulse brings the OPNs above the threshold for firing. Model simulations including this sequence of events recapitulate the results shown in Fig. 10.

Under the model, ipsilateral activation reduces the accumulated displacement signal maintained in the feedback integrator and produces a reacceleration during the deceleration phase of the saccade (Fig. 13D, red and blue traces). The latency of the reacceleration is attributed to a fixed, 5 ms delay in the feedback path, which is why the model produces the same latencies for early (red bars) and late (blue bar) activation pulses. This is an imperfect reflection of reality; the effect latency of ipsilateral OMV P-cell activation decreased when activation occurred later, experimentally (Table 1). This dependence of latency on pulse timing suggests a delay mechanism that depends on the real-time state of the saccades.

Delivering ipsi-and contralateral activating pulses to the model *simultaneously* produces saccades that mimic the mixed effects produced by activation of P-cells at the midline (compare Figs. 11 and 13E). An early contralateral activation pulse (red bar) reduces peak velocity, but the simultaneous ipsilateral activation pulse causes the model to over-compensate and produce hypermetria (Fig. 13E, red traces). Late, contralateral activation would normally truncate a fast saccade (Fig 13B, blue), but the simultaneous ipsilateral activation prevents this truncation (Fig. 13E, blue). We can understand this on the basis of DME and OPNs in the model. The ipsilateral activating pulse decreases the output of the feedback integrator, thereby increasing DME. This increase in DME balances the inhibitory effect of the simultaneous contralateral activating pulse, preventing the OPNs from being activated. The reacceleration after the pulse over-compensates for the perturbation, leading to hypermetria (Fig. 13E, blue traces). This is consistent with experimental data; saccades perturbed by midline stimulation were consistently hypermetric or normometric (Fig. 11).

In conclusion, manipulations to an established local feedback model suggest different roles for the contra-and ipsilateral OMV/cFN in saccade generation. The contralateral side contributes to the feedforward path, influencing the saccade at short latency, whereas the ipsilateral side contributes to the feedback path, integrating EBN activity and balancing the contralateral drive to allow the saccade to land on target.

## Discussion

Our results show that brief activation of OMV P-cells during a saccade changes its trajectory in midflight. The nature of the change depends on the direction of the saccade relative to the locus of stimulation. Stimulation decelerated contraversive saccades within ∼6 ms. If stimulation occurred near saccade onset, a compensatory reacceleration occurred during the deceleration phase, driving the saccades to land on or near the target. If stimulation occurred later, the compensatory reacceleration was velocity-dependent. Slower saccades reaccelerated and landed near the target. Faster saccades did not reaccelerate and were hypometric.

In contrast, OMV P-cell stimulation affected ipsiversive saccades at a longer delay. Under these circumstances, reacceleration effect during the saccade deceleration phase was not followed by a compensatory movement, so saccades were hypermetric. The amount of hypermetria grew with the duration of the activation pulse.

### Circuitry underlying the effects on contraversive saccades

The projections from OMV P-cells predict the effects of stimulation on contraversive saccades. OMV P-cells inhibit neurons in the cFN, which drive contralateral EBNs and IBNs. The EBNs, in turn, drive ipsilateral abducens motoneurons, and IBNs inhibit contralateral abducens motoneurons (Fig. 1A).

During optogenetic activation, contralateral P-cells inhibit the cFN of the same side, reducing its ability to excite EBNs and IBNs. Reduced activity in the EBNs and IBNs reduces the drive to the agonist abducens motoneurons and the inhibition of the antagonist abducens motoneurons, respectively. A short-latency slowing of the saccade is therefore expected. Given that the shortest latency of a simple spike following a light pulse is ∼3 ms (El-Shamayleh et al., 2017), and that optogenetic activation affected saccade acceleration after ∼6 ms, we conclude that P-cell simple spikes decelerate contraversive saccades within ∼3 ms.

Two additional observations support the idea that contralateral P-cell activation reduced the activity of the feedforward elements of a local feedback model (Robinson, 1975; Jürgens et al., 1981). First, T0 activation pulses slowed saccades but did not lead to hypometria. A reacceleration during the deceleration phase compensated for the attenuated acceleration phase. This effect is similar to those produced by brief OPN stimulation (Becker, 1981; Keller et al., 1996), and OPNs are known to inhibit neurons in the feedforward pathway (Langer and Kaneko, 1983; Strassman et al., 1987; Iwamoto et al., 2009).

Second, late activation pulses (T20 and T30), which occur during the saccade deceleration phase, truncated faster saccades. The local negative feedback model readily explains this phenomenon. The faster the saccade is, the lower the DME is when the activation pulse occurs. Low DME allows the stimulation-mediated reduction in EBN activity to disrupt the OPN latch, activating OPNs and terminating the saccade.

### Circuitry underlying the effects on ipsiversive saccades

In contrast to contralateral activation, ipsilateral activation produced hypermetric (non-compensated) saccades. The earliest activation pulses, even those as brief as 1.25 ms, elicited reacceleration >20 ms later. A similar effect can be produced by a brief inhibition of the feedback integrator in the model (Fig. 13).

The ipsilateral OMV/cFN complex may be part of the feedback integrator. The onset of the reacceleration effect produced by ipsilateral activation is similar to the time of the compensatory reacceleration produced by contralateral activation (Fig. 12), supporting the idea that a common mechanism underlies both phenomena. These effects are delayed relative to the pulse onset. We propose that the delay reflects delayed output from the feedback integrator (Fig. 13A). In this case, the contraversive compensatory reacceleration indicates the start of a closed-loop operation of the feedback circuit.

The ipsilateral OMV/cFN may receive input from the feedforward path to correct perturbations. Feedforward signals may be carried by the mossy fiber projection from the paramedian pontine reticular formation, which carries an efference copy of the EBN motor command (Yamada and Noda, 1987; Noda et al., 1990; Ohtsuka and Noda, 1992).

The left and right OMV likely contain the same microcircuits (Eccles, 1967). However, circuits on each side of the OMV apparently switch their role from a real-time feedforward controller during contraversive saccades to a feedback displacement integrator during ipsiversive saccades. As a feedforward controller, the OMV adjusts EBN and IBN activity to compensate for the eyes’ viscoelastic properties and to prevent untimely OPN reactivation that would otherwise truncate saccades. As a displacement integrator, the OMV determines the deceleration time course based on a running estimate of the displacement. Neurons outside the cerebellar cortex may influence this switching role and may lie in a feedback loop that, with OMV P-cells, forms a neural integrator. A cerebellar nucleocortical loop (Houck and Person, 2015) seems unlikely to form the integrator because the loop is not closed; tracer injections into the OMV label axon terminals in the fastigial nucleus densely, and some fastigial neurons retrogradely, but these populations do not overlap (Yamada and Noda, 1987). A more probable candidate is brainstem neurons that project directly or indirectly to the OMV as mossy fibers. If these mossy fibers play this role, we predict that they should exhibit a gradual increase of activity just before and during a saccade like a long-lead burst neuron, and their activity should be different during contra-and ipsiversive saccades (i.e., they should have directional tuning, Ohtsuka and Noda, 1992; Prsa et al., 2009). The involvement of long-lead burst neurons in a displacement integrator has been predicted previously (Scudder, 1988).

### Comparison with cFN inactivation studies

Activation near the midline of the vermis, which presumably inhibited both left and right cFNs, produced saccades that were slightly hypermetric and slow, with a prolonged deceleration phase. These effects are similar to those produced by bilateral cFN inactivation (Robinson et al., 1993).

Both long-lasting ipsilateral cFN inactivation and our brief ipsilateral P-cell activation produced hypermetria. Common mechanisms may underlie both effects, but the literature is not entirely consistent with this idea. In one study, ipsilateral cFN inactivation produced hypermetria consistent with our results, i.e., no change in the saccade acceleration phase or peak velocity but a prolonged saccade deceleration phase (Goffart et al., 2004). However, in an earlier study, saccade hypermetria was associated with increased peak velocity and saccade duration (Robinson et al., 1993). The reason for this discrepancy is unclear.

Contralateral cFN inactivation slows saccades and produces hypometria (Robinson et al., 1993; Goffart et al., 2004). Similarly, our early, brief P-cell activation also slowed saccades, but they were nearly normometric due to compensatory reacceleration. We speculate that following contralateral cFN inactivation, the intact ipsilateral cFN compensates for the resultant hypometria but incompletely. This partial compensation may account for the smaller hypometric deficit of contraversive saccades after unilateral cFN inactivation (Iwamoto and Yoshida, 2002).

### Correlation with P-cell simple spike responses during saccades

Activating OMV P-cells optogenetically increases simple spike (SS) firing rates (El-Shamayleh et al., 2017) and alters saccade trajectories. In the absence of experimental manipulation, contralateral P-cells exhibit bursts of SS activity during saccade acceleration, and they pause during saccade deceleration (Herzfeld et al., 2015, their Fig. 4, green; Herzfeld and Shadmehr, 2016). The duration of the pause may be critically important for sustaining saccade deceleration. During the deceleration phase of normal, visually guided saccades, the pause in contralateral SS activity is briefer and shallower during fast saccades than during slow ones (Herzfeld et al., 2015, their Fig. 3f). The prolonged pause during slow saccades may allow the deceleration phase to continue long enough for the eyes to reach the target. In our T20 and T30 conditions, we stimulated contralateral P-cells during the deceleration phase, filling the pause with optogenetically produced SS. These abbreviated pauses may be why fast saccades were truncated and slow saccades were not.

The ipsilateral P-cell population fires smaller saccade-related bursts than the contralateral P-cell population (Herzfeld et al., 2015, their Fig. 4, brown). Ipsilateral P-cells do not pause during saccade deceleration but continue to fire for ∼50 ms after the saccade termination. The functional significance of this activity is unclear, but its post-saccade persistence is consistent with the passive discharge of a leaky integrator (Kustov and Robinson, 1995; Nichols and Sparks, 1995). In future experiments, it will be important to analyze the interaction between saccade-related and optogenetically produced spiking activity in P-cells.

## Conclusion

In conclusion, contralateral and ipsilateral OMV P-cells play distinct roles in controlling saccades. These roles can be understood within the framework of an established local negative feedback model. The contralateral OMV likely houses the feedforward inverse model of the oculomotor mechanics to estimate the appropriate saccade motor command, as evidenced by the low-latency deceleration effect on saccade trajectory and the compensatory reacceleration observed following perturbations. Conversely, the ipsilateral OMV, which produces an uncompensated long-latency reacceleration effect, is likely involved in the feedback forward model that predicts eye displacement. This prediction is likely accomplished by temporal integration of the efference copy of the saccade motor command. This possibility could be tested by briefly inhibiting P-cells. Under these conditions we would expect faster but normometric contraversive saccades and hypometric ipsiversive saccades.

## Acknowledgments

National Institute of Health fundings: EY028902 (RS), EY030441 (GH), OD010425, RR00166 (Washington National Primate Research Center), P30EY001730 (Vision Research Core of University of Washington). The authors thank Dr. Albert Fuchs for his valuable comments and editorial assistance and Shane Gibson and Yasmine El-Shamayleh for assistance with histology.

The authors declare no competing financial interests.

## References

Bahill AT, Clark MR, Stark L (1975) The main sequence, a tool for studying human eye movements. Mathematical Biosciences 24:191–204.

Barash S, Melikyan A, Sivakov A, Zhang M, Glickstein M, Thier P (1999) Saccadic dysmetria and adaptation after lesions of the cerebellar cortex. The Journal of neuroscience : the official journal of the Society for Neuroscience 19:10931–10939.

Barton EJ, Nelson JS, Gandhi NJ, Sparks DL (2003) Effects of partial lidocaine inactivation of the paramedian pontine reticular formation on saccades of macaques. Journal of neurophysiology 90:372–386.

Becker W, King, W.M., Fuchs, A.F., Jürgens, R., Johanson, G., Kornhuber, H.H. (1981) Accuracy of goal-directed saccades and mechanisms of error correction. In: Progress in Oculomotor Research (Fuchs AF, Becker, W., ed), pp 29-37. New York: Elsevier.

Collins CC, O’Meara D, Scott AB (1975) Muscle tension during unrestrained human eye movements. The Journal of physiology 245:351–369.

Crandall WF, Keller EL (1985) Visual and oculomotor signals in nucleus reticularis tegmenti pontis in alert monkey. Journal of neurophysiology 54:1326–1345.

Eccles JC (1967) Circuits in the cerebellar control of movement. Proceedings of the National Academy of Sciences of the United States of America 58:336–343.

El-Shamayleh Y, Kojima Y, Soetedjo R, Horwitz GD (2017) Selective Optogenetic Control of Purkinje Cells in Monkey Cerebellum. Neuron 95:51–62.e54.

Fuchs AF (1967) Saccadic and smooth pursuit eye movements in the monkey. The Journal of physiology 191:609–631.

Fuchs AF, Luschei ES (1971) Development of isometric tension in simian extraocular muscle. The Journal of physiology 219:155–166.

Fuchs AF, Kaneko CR, Scudder CA (1985) Brainstem control of saccadic eye movements. Annual review of neuroscience 8:307–337.

Fuchs AF, Scudder CA, Kaneko CR (1988) Discharge patterns and recruitment order of identified motoneurons and internuclear neurons in the monkey abducens nucleus. Journal of neurophysiology 60:1874–1895.

Fujikado T, Noda H (1987) Saccadic eye movements evoked by microstimulation of lobule VII of the cerebellar vermis of macaque monkeys. The Journal of physiology 394:573–594.

Goffart L, Chen LL, Sparks DL (2004) Deficits in saccades and fixation during muscimol inactivation of the caudal fastigial nucleus in the rhesus monkey. Journal of neurophysiology 92:3351–3367.

Goldstein H, Robinson D (1984) A two-element oculomotor plant model resolves problems inherent in a single-element plant model. In: Soc Neurosci Abstr, p 909.

Harris CM, Wolpert DM (2006) The main sequence of saccades optimizes speed-accuracy trade-off. Biological cybernetics 95:21–29.

Herzfeld DJ, Shadmehr R (2016) Cerebellar output encodes a corrective saccadic command (Commentary on Sun et al.). The European journal of neuroscience 44:2528–2530.

Herzfeld DJ, Kojima Y, Soetedjo R, Shadmehr R (2015) Encoding of action by the Purkinje cells of the cerebellum. Nature 526:439–442.

Houck BD, Person AL (2015) Cerebellar Premotor Output Neurons Collateralize to Innervate the Cerebellar Cortex. The Journal of comparative neurology 523:2254–2271.

Iwamoto Y, Yoshida K (2002) Saccadic dysmetria following inactivation of the primate fastigial oculomotor region. Neuroscience letters 325:211–215.

Iwamoto Y, Kaneko H, Yoshida K, Shimazu H (2009) Role of glycinergic inhibition in shaping activity of saccadic burst neurons. Journal of neurophysiology 101:3063–3074.

Judge SJ, Richmond BJ, Chu FC (1980) Implantation of magnetic search coils for measurement of eye position: an improved method. Vision research 20:535–538.

Jürgens R, Becker W, Kornhuber HH (1981) Natural and drug-induced variations of velocity and duration of human saccadic eye movements: evidence for a control of the neural pulse generator by local feedback. Biological cybernetics 39:87–96.

Kaneko CR (1997) Eye movement deficits after ibotenic acid lesions of the nucleus prepositus hypoglossi in monkeys. I. Saccades and fixation. Journal of neurophysiology 78:1753–1768.

Keller EL, Gandhi NJ, Shieh JM (1996) Endpoint accuracy in saccades interrupted by stimulation in the omnipause region in monkey. Visual neuroscience 13:1059–1067.

Kojima Y, Soetedjo R, Fuchs AF (2010) Changes in simple spike activity of some Purkinje cells in the oculomotor vermis during saccade adaptation are appropriate to participate in motor learning. The Journal of neuroscience : the official journal of the Society for Neuroscience 30:3715–3727.

Kojima Y, Robinson FR, Soetedjo R (2014) Cerebellar fastigial nucleus influence on ipsilateral abducens activity during saccades. Journal of neurophysiology 111:1553–1563.

Kojima Y, Ting JT, Soetedjo R, Gibson SD, Horwitz GD (2021) Injections of AAV Vectors for Optogenetics in Anesthetized and Awake Behaving Non-Human Primate Brain. Journal of visualized experiments : JoVE.

Krauzlis RJ, Miles FA (1998) Role of the oculomotor vermis in generating pursuit and saccades: effects of microstimulation. Journal of neurophysiology 80:2046–2062.

Kustov AA, Robinson DL (1995) Modified saccades evoked by stimulation of the macaque superior colliculus account for properties of the resettable integrator. Journal of neurophysiology 73:1724–1728.

Langer TP, Kaneko CR (1983) Efferent projections of the cat oculomotor reticular omnipause neuron region: an autoradiographic study. The Journal of comparative neurology 217:288–306.

Nichols MJ, Sparks DL (1995) Nonstationary properties of the saccadic system: new constraints on models of saccadic control. Journal of neurophysiology 73:431–435.

Noda H, Sugita S, Ikeda Y (1990) Afferent and efferent connections of the oculomotor region of the fastigial nucleus in the macaque monkey. The Journal of comparative neurology 302:330–348.

Noda H, Murakami S, Warabi T (1991) Effects of fastigial stimulation upon visually-directed saccades in macaque monkeys. Neuroscience research 10:188–199.

Noda H, Murakami S, Yamada J, Tamada J, Tamaki Y, Aso T (1988) Saccadic eye movements evoked by microstimulation of the fastigial nucleus of macaque monkeys. Journal of neurophysiology 60:1036–1052.

Ohtsuka K, Noda H (1992) Burst discharges of mossy fibers in the oculomotor vermis of macaque monkeys during saccadic eye movements. Neuroscience research 15:102–114.

Optican LM, Miles FA (1985) Visually induced adaptive changes in primate saccadic oculomotor control signals. Journal of neurophysiology 54:940–958.

Prsa M, Dash S, Catz N, Dicke PW, Thier P (2009) Characteristics of responses of Golgi cells and mossy fibers to eye saccades and saccadic adaptation recorded from the posterior vermis of the cerebellum. The Journal of neuroscience : the official journal of the Society for Neuroscience 29:250–262.

Robinson DA (1964) The mechanics of human saccadic eye movement. The Journal of physiology 174:245–264.

Robinson DA (1975) Oculomotor control signals. In: Basic mechanisms of ocular motility and their clinical implications (P. Bach-y-Rita GL, ed), pp 337-374. Oxford: Pergamon.

Robinson FR, Straube A, Fuchs AF (1993) Role of the caudal fastigial nucleus in saccade generation. II. Effects of muscimol inactivation. Journal of neurophysiology 70:1741–1758.

Sasaki S, Shimazu H (1981) Reticulovestibular organization participating in generation of horizontal fast eye movement. Annals of the New York Academy of Sciences 374:130–143.

Scudder CA (1988) A new local feedback model of the saccadic burst generator. Journal of neurophysiology 59:1455–1475.

Scudder CA, Fuchs AF, Langer TP (1988) Characteristics and functional identification of saccadic inhibitory burst neurons in the alert monkey. Journal of neurophysiology 59:1430–1454.

Soetedjo R, Fuchs AF (2006) Complex spike activity of purkinje cells in the oculomotor vermis during behavioral adaptation of monkey saccades. The Journal of neuroscience : the official journal of the Society for Neuroscience 26:7741–7755.

Soetedjo R, Kojima Y (2022) Optogenetics in Complex Model Systems (Non-Human Primate). In: Measuring Cerebellar Function (Sillitoe RV, ed), pp 305–321. New York, NY: Springer US.

Soetedjo R, Kaneko CR, Fuchs AF (2002) Evidence that the superior colliculus participates in the feedback control of saccadic eye movements. Journal of neurophysiology 87:679–695.

Soetedjo R, Kojima Y, Fuchs AF (2008) Complex spike activity in the oculomotor vermis of the cerebellum: a vectorial error signal for saccade motor learning? Journal of neurophysiology 100:1949–1966.

Soetedjo R, Kojima Y, Fuchs AF (2019) How cerebellar motor learning keeps saccades accurate. Journal of neurophysiology 121:2153–2162.

Sparks D, Rohrer WH, Zhang Y (2000) The role of the superior colliculus in saccade initiation: a study of express saccades and the gap effect. Vision research 40:2763–2777.

Strassman A, Highstein SM, McCrea RA (1986) Anatomy and physiology of saccadic burst neurons in the alert squirrel monkey. I. Excitatory burst neurons. The Journal of comparative neurology 249:337–357.

Strassman A, Evinger C, McCrea RA, Baker RG, Highstein SM (1987) Anatomy and physiology of intracellularly labelled omnipause neurons in the cat and squirrel monkey. Experimental brain research 67:436–440.

Sylvestre PA, Cullen KE (1999) Quantitative analysis of abducens neuron discharge dynamics during saccadic and slow eye movements. Journal of neurophysiology 82:2612–2632.

Takagi M, Zee DS, Tamargo RJ (1998) Effects of lesions of the oculomotor vermis on eye movements in primate: saccades. Journal of neurophysiology 80:1911–1931.

Takahashi M, Sugiuchi Y, Na J, Shinoda Y (2022) Brainstem Circuits Triggering Saccades and Fixation. The Journal of Neuroscience 42:789–803.

Taylor JR (1997) An introduction to error analysis : the study of uncertainties in physical measurements, 2nd ed Edition. [Place of publication not identified]: University Science.

van Gisbergen JAM, Robinson DA, Gielen S (1981) A quantitative analysis of generation of saccadic eye movements by burst neurons. Journal of neurophysiology 45:417–442.

Wilcox RR (2022) Chapter 5 - Comparing Two Groups. In: Introduction to Robust Estimation and Hypothesis Testing (Fifth Edition) (Wilcox RR, ed), pp 153-251: Academic Press.

Xu-Wilson M, Tian J, Shadmehr R, Zee DS (2011) TMS perturbs saccade trajectories and unmasks an internal feedback controller for saccades. The Journal of neuroscience : the official journal of the Society for Neuroscience 31:11537–11546.

Yamada J, Noda H (1987) Afferent and efferent connections of the oculomotor cerebellar vermis in the macaque monkey. The Journal of comparative neurology 265:224–241.

